# The ribosome-associated quality control factor Vms1 protects mitochondrial import and homeostasis during translation stress

**DOI:** 10.64898/2026.07.22.740037

**Authors:** Gülsah Göktas, Vitasta Tiku, Nikita A. Kvasov, Andrei Makhon, Michal Breker, Fabian den Brave, Markus Raeschle, Zuzana Storchova, Yury S. Bykov

## Abstract

Mitochondrial protein synthesis and import are tightly coordinated to maintain cellular proteostasis, yet how cytosolic translation stress affects mitochondrial homeostasis remains poorly understood. Here, we investigated the cellular consequences of general translation stress using low-dose translation inhibitors in *Saccharomyces cerevisiae*. Genome-wide phenotypic screening revealed that the deletion of tRNA-hydrolase *VMS1* involved in mitochondria-associated ribosomal quality control (mitoRQC) causes a unique hypersensitivity to low-dose cycloheximide. Quantitative proteomics demonstrated that translation stress triggers specific depletion of mitochondrial proteins, reduced respiratory capacity, and impaired mitochondrial membrane potential. Most notably, cytosolic translation stress strongly inhibited mitochondrial protein import, an effect that was substantially exacerbated in *vms1Δ* cells. We further identify the mitochondrial AAA-ATPase Msp1 as a protective factor during translation stress. Msp1 expression increased in *vms1Δ* cells, and its overexpression restored growth under stress, linking co-translational quality control to mitochondrial protein import surveillance. Together, our findings reveal mitochondrial protein import as a major target of cytosolic translation stress and uncover functional cooperation between mitoRQC and Msp1 in safeguarding mitochondrial proteostasis and cell survival.

## Introduction

Protein synthesis is one of the most ancient cellular processes (Fer et al., 2025). It is also relatively slow, energy demanding and error-prone (Tanenbaum et al., 2014; Metzl-Raz et al., 2017; Schuller and Green, 2018) and its decline in efficiency is one of the ageing hallmarks (Di Fraia et al., 2025). Different factors such as damaged mRNA, amino acid or tRNA absence, xenobiotics, oxidative stress, or specific codon sequences cause errors and delays in translation elongation and trigger translation stress. (Schuller and Green, 2018; Wu et al., 2020; Yan et al., 2019). One of the key events that activates cellular responses to translation stress is ribosome stalling that results in ribosome collisions. The collided ribosomes are recognized by conserved ribosome rescue systems and stress signaling pathway components (Matsuo et al., 2017; Simms et al., 2017; Juszkiewicz et al., 2018; Ikeuchi et al., 2019; Wu et al., 2020; Pochopien et al., 2021; Zhou et al., 2025). Translation rescue systems can cleave the mRNAs trapped in stalled ribosomes and split ribosome subunits (Chiabudini et al., 2014; Guydosh and Green, 2014; Shcherbik et al., 2016; D’Orazio et al., 2019). The resulting large subunits (60S) that still contain tRNA-bound nascent chains are recycled by the ribosome-associated quality control (RQC) pathway (reviewed in (Filbeck et al., 2022) and (Inada, 2026)). The 60S-bound nascent chain is ubiquitinated and targeted to degradation (Bengtson and Joazeiro, 2010; Li et al., 2025a). To promote efficient ubiquitination and degradation, nascent chain is also extended by addition of alanine and threonine residues creating C-terminal extensions called CAT-tails (Shen et al., 2015; Kostova et al., 2017; Sitron and Brandman, 2019; Ennis et al., 2025). Finally, the nascent chain is released from the large subunit by tRNA hydrolases and targeted to the proteasome (Brandman et al., 2012; Kuroha et al., 2018; Verma et al., 2018; Zurita Rendón et al., 2018; Su et al., 2019). Together, these mechanisms help to maintain translation fidelity and cell viability under a wide range of stress conditions.

While the molecular mechanisms of stalled ribosome rescue and recycling are well understood, the physiological consequences of translation stress and RQC failure are only beginning to be uncovered. Ribosome stalling and the defects of the RQC components are associated with neurological and autoimmune disorders (Chu et al., 2009; Ishimura et al., 2014; van Haaften-Visser et al., 2017; Martin et al., 2020). The cellular mechanisms behind the physiological effects of translation stress were linked with proteostasis failure caused by the accumulation of CAT-tailed proteins (Choe et al., 2016). Such proteins can create toxic aggregates and deplete the cytosolic chaperone pool (Defenouillère et al., 2013; Yonashiro et al., 2016; Choe et al., 2016; Sitron et al., 2020). The toxicity of CAT-tailed proteins depends on their composition and the activity of the other RQC components (Choe et al., 2016; Sitron et al., 2020; Udagawa et al., 2021). It remains to be determined how CAT tails contribute to the effects of translation stress in different cell types and organisms. The effects of translation stress are not limited to cytosolic proteostasis. Overexpression of ER- and mitochondria-targeted non-stop (NS) proteins (from mRNAs lacking stop codons) showed that ribosome rescue and RQC pathways are required to maintain the function of these organelles (Izawa et al., 2012; Izawa et al., 2017; Bertram et al., 2025). The studies in yeast identified tRNA hydrolase Vms1 (homolog of human ANKZF1) as an important factor linking cytosolic RQC and mitochondrial homeostasis (Bertram et al., 2025; Izawa et al., 2017). Vms1 prevents excessive CAT-tailing of aberrant translation products destined to the mitochondrial matrix where they create toxic aggregates and inhibit respiration. Thus, the effects of cytosolic translation stress can be far reaching and can affect mitochondrial homeostasis.

The majority of mitochondrial proteins are synthesized by cytosolic ribosomes and imported via specialized translocation pathways (Neupert and Herrmann, 2007). This dependence on protein import creates tight connections between mitochondrial function and cytosolic proteostasis. First, the import requires cytosolic chaperones to deliver unfolded precursor proteins (precursors) to the translocase of the outer membrane (TOM) complex (Young et al., 2003; Zara et al., 2009; Hoseini et al., 2016; Opaliński et al., 2018; Juszkiewicz et al., 2025). Second, mitochondrial dysfunction or specific translocase inhibition causes cytosolic accumulation and aggregation of unfolded precursors, and induces cytosolic stress response pathways such as the integrated stress response (ISR), heat shock response, and mitochondrial import surveillance pathway MitoCPR (Boos et al., 2019; Fessler et al., 2020; Guo et al., 2020; Wang and Chen, 2015; Weidberg and Amon, 2018; Wrobel et al., 2015). Finally, both induced, and constitutive mechanisms such as mitoCPR and mitochondrial-import associated degradation (mitoTAD) directly surveil the TOM complex. These pathways can detect precursors arrested in the TOM complex and promote their extraction and degradation by the ubiquitin-proteasome system (Kim et al., 2024; Mårtensson et al., 2019; Schulte et al., 2023). In summary, mitochondrial dysfunction can have strong effects on cytosolic proteostasis and stress signaling. The mechanisms that counteract such stresses and maintain mitochondrial homeostasis rely heavily on post-translational quality control pathways such as chaperone networks and the ubiquitin-proteasome system (UPS). The connections between mitochondrial function and co-translational quality control pathways are much less understood and rely on studies that used overexpression of individual model NS proteins.

Here we use low dose translation inhibitor treatment to induce general cytosolic translation stress in *Saccharomyces cerevisiae* (hereafter called yeast) and investigate its effects on cellular and mitochondrial homeostasis. Using a genetic screen, we identify Vms1 as a major determinant of cellular viability under these conditions. We use proteomics and functional assays to demonstrate that cytosolic translation stress broadly affects mitochondrial homeostasis identifying a strong link between co-translational quality control and mitochondrial function. We also show that mitochondrial protein import is strongly inhibited in *vms1Δ* mutant under translation stress conditions highlighting important role of this protein in supporting cell viability. Finally, we find that mitoCPR factor Msp1 can rescue the detrimental effects of translation stress in *vms1Δ* mutant, connecting co- and post-translational mechanisms that maintain mitochondrial homeostasis.

## Results

### VMS1 deletion increases sensitivity to prolonged mild translation stress

To understand the role of ribosome rescue and RQC components in cell survival under general translation stress we selected several mutant strains to compare their response to different concentrations of translation inhibitor cycloheximide (CHX) using a drop dilution assay. We chose to test the RQC factors Ltn1, Rqc2, Vms1, the ribosome collision sensor Hel2, and splitting factor Dom34 that act at different stages of stalled ribosome rescue and recycling (Fig. 1A)(Bengtson and Joazeiro, 2010; van den Elzen et al., 2014; Shen et al., 2015; Verma et al., 2018; Ikeuchi et al., 2019). The growth of all the mutants was impaired on glucose (fermentative) and glycerol-containing (respiratory) media at the CHX concentration of 100 ng/ml while the WT strain grew normally (Fig 1B). The reduction of CHX concentration to 10 ng/ml rescued the growth of all the mutants except the *vms1Δ* on both media types. On fermentative media with the minimal CHX concentration *vms1Δ* strain grew only slightly slower than WT, while on respiratory media the growth reduction relative to the WT strain was much more prominent, consistent with the role of Vms1 in protecting mitochondria from ribotoxic stress (Izawa et al., 2017). Testing the growth of the RQC mutants in the presence of a different translation inhibitor (anisomycin) and the mRNA damaging agent methyl methanesulfonate (MMS) also showed stronger sensitivity of the *vms1Δ* strain compared to the others (Fig. S1A, B). The *vms1Δ* strain growth on low CHX media was fully rescued by a plasmid encoding WT Vms1 expressed under its endogenous promoter (Fig. S1C). The Vms1 R288A mutant that cannot hydrolase tRNAs enabled partial rescue on fermentative media but not on respiratory media, while Vms1Δ*VIM* lacking the Cdc48-interacting sequence showed partial rescue on both medias (Fig. S1C), similarly to the data reported previously (Zurita Rendón et al., 2018). The growth reduction observed by drop dilutions could result from cell death or from regulatory growth inhibition. To distinguish these two possibilities, we performed short-term CHX treatment of WT and mutant cell cultures after which the cells were plated on rich media without CHX to count the number of colony-forming units. We found that *vms1Δ* cells significantly lost viability already after 2 h of CHX treatment indicating cell death (Fig. 1C). The hypersensitivity of the *VMS1* deletion strains to treatments that interrupt translation indicates a special role for this protein in protecting cellular homeostasis under ribotoxic stress.

**Figure 1.**
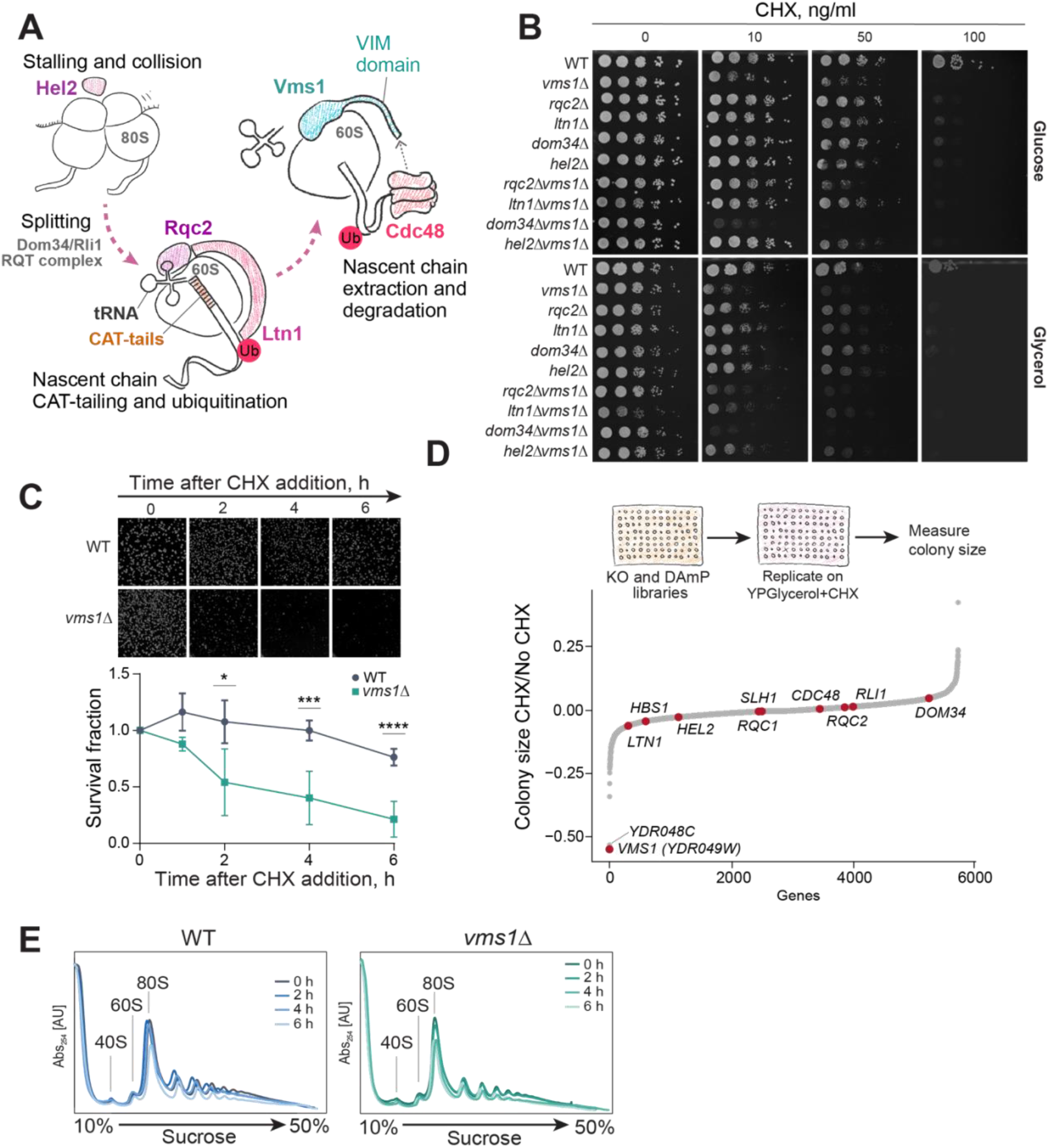
The deletion of VMS1 results in hypersensitivity to low dose translation inhibitors. **(A)** Overview of stalled-ribosome rescue and ribosome-associated quality control. Partial inhibition of translation promotes ribosome collisions, which are recognized by the E3 ubiquitin ligase Hel2. Stalled ribosomes are dissociated by the Dom34-Hbs1-Rli1 rescue system or RQT complex, generating a 60S subunit carrying the nascent chain linked to tRNA. Rqc2 binds the resulting 60S-peptidyl-tRNA complex and promotes the C-terminal addition of alanine and threonine residues (CAT-tailing), while Ltn1 ubiquitylates the nascent chain. Vms1 hydrolyzes the peptidyl-tRNA linkage and cooperates with Cdc48 in nascent-chain release. **(B)** Serial-dilution growth assay of the indicated single and double mutants on the indicated carbon sources and CHX concentrations. Plates were incubated at 30°C. **(C)** Survival of WT and *vms1Δ* cells after treatment with 100 ng/ml CHX for 1 to 6 h. Colony formation was assessed after 2 days and normalized to untreated cells. Data represents three independent experiments. Statistical significance was determined by two-way ANOVA followed by Šídák’s multiple-comparisons test. *P < 0.05, **P < 0.001, ***P < 0.0001. **(D)** Genome-wide screening of the non-essential gene deletion and essential-gene DAmP collections on YPGlycerol containing 100 ng/ml CHX. Colony sizes were quantified. **(E)** Polysome profiles of WT and *vms1*Δ cells treated with 100 ng/ml CHX for 2–6 h. Lysates were separated on 10–50% sucrose gradients, and RNA absorbance was recorded during fractionation.

Next, we decided to test if any other yeast gene deletion has a hypersensitive phenotype similar to *vms1Δ*. For this, we plated the arrayed collection of yeast non-essential deletion mutants (Giaever et al., 2002) and the collection of essential gene mutants constructed using the decreased abundance by mRNA perturbation (DAmP) approach (Breslow et al., 2008) on solid respiratory media with low CHX concentration, and measured the reduction of colony size relative to media without CHX. Surprisingly, most strains showed none, or a very mild growth defect, while only the deletions of *VMS1* (*YDR049W*) and an overlapping dubious open reading frame *YDR048C* had a strong growth reduction (Fig. 1D, Table S1). The deletion of *YDR048C* removes the promoter and the first 150 bp of the *VMS1* gene and also creates a non-functional *vms1* allele. This result suggests that among the tested genes *VMS1* plays a unique role in protection against ribotoxic stress.

To understand possible redundancies within the RQC pathway, we tested genetic interactions of *VMS1* with RQC components whose single deletions did not result in strong phenotypes. We found that the additional deletion of *DOM34* exacerbated CHX sensitivity, resulting in an additive phenotype (Fig. 1B). This suggests that Dom34 and Vms1 make at least partially distinct contributions to ribosome rescue and that Dom34 may support alternative recycling pathways that do not depend on Vms1 (Inada, 2026). In contrast, deletion of *LTN1* alleviated the phenotype, whereas deletion of *RQC2* resulted in a weaker rescue (Fig. 1B), consistent with a previous report (Zurita Rendón et al., 2018). While the effect of *RQC2* deletion may be explained by the lack of CAT-tailing, the rescue mechanism behind the *LTN1* deletion is unknown (Izawa et al., 2017; Zurita Rendón et al., 2018). We additionally verified that upon CHX treatment the *vms1Δ* strain indeed accumulated high molecular weight ubiquitinated species that were absent when *LTN1* was deleted (Fig. S1D). We also find that on respiratory media without CHX the double deletion *vms1Δltn1Δ* is synthetically sick. This phenotype was explained by the increased import of CAT-tailed proteins into the mitochondria because the cytosolic degradation of such nascent chain is inhibited (Bertram et al., 2025; Izawa et al., 2017). Interestingly, CHX addition reverses this negative genetic interaction of *VMS1* and *LTN1* on respiratory media (Fig. 2B). This suggests that unspecific translation stress caused by CHX may lead to a phenotype that is distinct from the phenotype caused by CAT-tailed peptide accumulation in the mitochondrial matrix that only happens in *vms1Δltn1Δ* strain on respiratory media.

**Figure 2.**
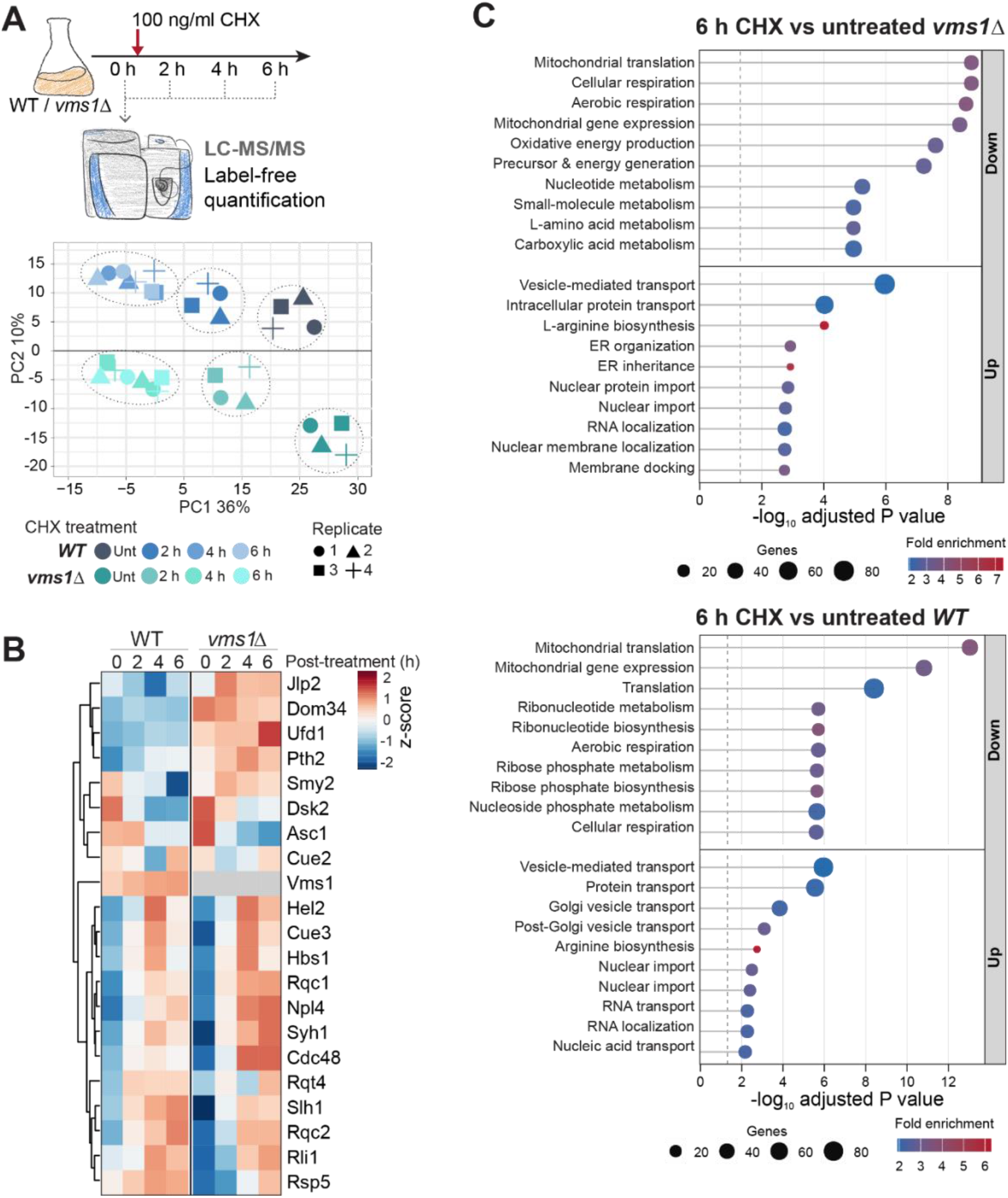
Proteome-level response of WT and *vms1Δ* cells to cycloheximide treatment. **(A)** Experimental workflow and principal component analysis (PCA). WT and *vms1*Δ cells were grown in galactose-containing medium and treated with 100 ng/ml CHX for 2, 4, or 6 h; untreated cells served as the 0 h control. Four biological replicates were analyzed per genotype and condition by label-free LC–MS/MS. Colors indicate genotype and treatment duration, and symbols represent individual replicates. **(B)** Heatmap of the relative abundance of RQC-related proteins in WT and *vms1Δ* cells during CHX treatment. Proteins were hierarchically clustered according to their abundance profiles. **(C)** GO enrichment analysis of proteins significantly increased or decreased after 6 h of CHX treatment relative to the corresponding untreated condition in *vms1Δ* (top) and WT (bottom) cells. Dot size indicates gene count, color indicates fold enrichment, and the x-axis shows −log10(adjusted P value).

Considering that Vms1 has a broader nascent chain release function in the RQC pathway, we checked how translation is affected under low CHX treatment conditions. Using a short pulse-chase assay for radiolabeled methionine incorporation we show that the prior treatment with low CHX for 2 to 6 h decreases overall protein synthesis, but this happens similarly in WT and *vms1Δ* cells (Figure S1D). We also checked if the ribotoxic stress and deletion of *VMS1* affected the translation status by performing polysome fractionation in sucrose density gradients. We find that there is no accumulation of 60S neither in WT nor in the mutant and only slight decrease of the 80S and polysome peaks upon CHX treatment in both strains (Fig 1E, Fig. S1E). We hence conclude that Vms1 depletion has an effect on cell survival under ribotoxic stress that cannot be completely explained by a general defect in translation or accumulation of non-recycled ribosome subunits.

### Proteomics reveals broad changes in translation, metabolism, and protein transport during ribotoxic stress

To investigate the cellular consequences of ribotoxic stress we treated the cultures of WT and *vms1Δ* strains with 100 ng/ml CHX for 2, 4, and 6 h and analyzed the cellular protein content with liquid chromatography-coupled tandem mass spectrometry (LC-MS/MS)(Fig. 2A) We detected at least 3000 proteins common for all samples (Fig. S2A, Table S2). Principle component analysis of the label free quantification (LfQ) intensities revealed that the CHX treatment induces strong proteome changes in both WT and mutant cells that are visualized along the primary component 1 (PC1) axis, while the differences between mutant and WT are more smaller and result in the variance along the PC2 axis (Fig 2A). Based on LfQ intensities, we calculated fold changes (FCs) in protein abundance between untreated and 6 h-treated samples within each strain (WT and *vms1Δ*), as well as between WT and *vms1Δ* strains at both the 0 h, and 6 h time points. Consistent with PCA analysis, only a few proteins had significant FC differences between WT and *vms1Δ* strain while hundreds were significantly changed upon CHX treatment (Fig S2B, Table S2).

We first checked how the abundance of RQC components and ribosome rescue factors changes upon CHX treatment. In line with an important role of these proteins in surviving translation stress, most of them were upregulated upon CHX treatment both in WT and *vms1Δ* (Fig. 2B). Interestingly, the few proteins, including Jlp2 and Dom34 are upregulated only in the mutant. Jlp2 was recently identified as an alternative factor that promotes tRNA hydrolysis to release nascent chains from stalled ribosomes (Iyer et al., 2025). Specific upregulation of Jlp2 in *vms1Δ* is consistent with its role in an alternative rescue pathway. Additional deletion of Dom34 made the *vms1Δ* growth phenotype stronger (Fig. 1C), further supporting the hypothesis that it may also be a part of ribosome recycling pathway that is alternative to the pathway mediated by Vms1 (Fig. 1B). This result points to the possibility that different splitting mechanisms such as either Dom34/Rli1 or the one based on RQT complex may culminate in the 60S-tRNA-nascent chain complexes that require different tRNA hydrolysis factors (Inada, 2026). Overall, the clear effect of CHX treatment on ribosome rescue factor abundance demonstrates the validity of our proteomics data. However, these factors only represent a small fraction of the proteome changes observed under translation stress.

To understand which functional protein categories drive the major proteome changes upon CHX treatment we performed gene ontology (GO) enrichment analysis using ShinyGO (Ge et al., 2020). Since translation stress often results in metabolic remodeling to supply more amino acids and in inhibition of ribosome biogenesis we expected to find changes in these functional categories (Yan and Zaher, 2021). Indeed, “Arginine biosynthesis” was among the upregulated GO terms in both WT and *vms1Δ* mutant (Fig. 2C). More detailed examination of individual amino acid biosynthesis pathways showed that many of the arginine, lysine and histidine biosynthesis enzymes increased their abundance after 2-4 h of CHX treatment, and this effect is more pronounced in the *vms1Δ* strain (Fig. S3). Interestingly, the most prominent GO terms downregulated upon CHX treatment included “Mitochondrial translation” and “Cellular respiration” (Fig. 2C). Comparison of cytosolic and mitochondrial translation components revealed a distinct trend toward altered abundance of mitoribosomal proteins, whereas cytosolic ribosomal proteins showed no comparable pattern. (Fig. S2C). To sum up, our proteomic analysis uncovered that cytosolic translation stress induces metabolic adaptation, which is consistent with the previous reports, but also strongly affects the mitochondrial proteome suggesting a tight link between translational quality control in the cytosol and mitochondrial functions.

### Respiration and mitochondrial import are impaired under cytosolic translation stress

Next, we aimed to characterize the effect of cytosolic translation stress and *VMS1* deletion on mitochondrial proteome and functions in more detail. Among proteins associated with respiration and metabolism, selected OXPHOS subunits and TCA cycle enzymes tended to be more abundant in *vms1*Δ cells without CHX treatment, however there were no differences that were similar for many proteins. (Fig. 3A, Fig. S4A, Table S2). Together with the upregulation of amino acid biosynthesis pathways such remodeling of the mitochondrial proteome might be a metabolic adaptation to survive translation stress. For example, in rapidly growing cancer cells mitochondria are not essential to produce energy, but the correct function of OXPHOS is necessary to produce cellular building blocks (Zhu et al., 2026). We then checked if mitochondrial function is essential to survive translation stress under non-respiratory conditions by treating WT and *vms1Δ* cells with a combination of low level of CHX and either Complex IV inhibitor antimycin A, ATP-synthase inhibitor oligomycin, or protonophore CCCP. The addition of these compounds did not worsen the phenotypes caused by CHX, suggesting that respiratory functions are not specifically required for stress survival (Fig. S4B).

**Figure 3.**
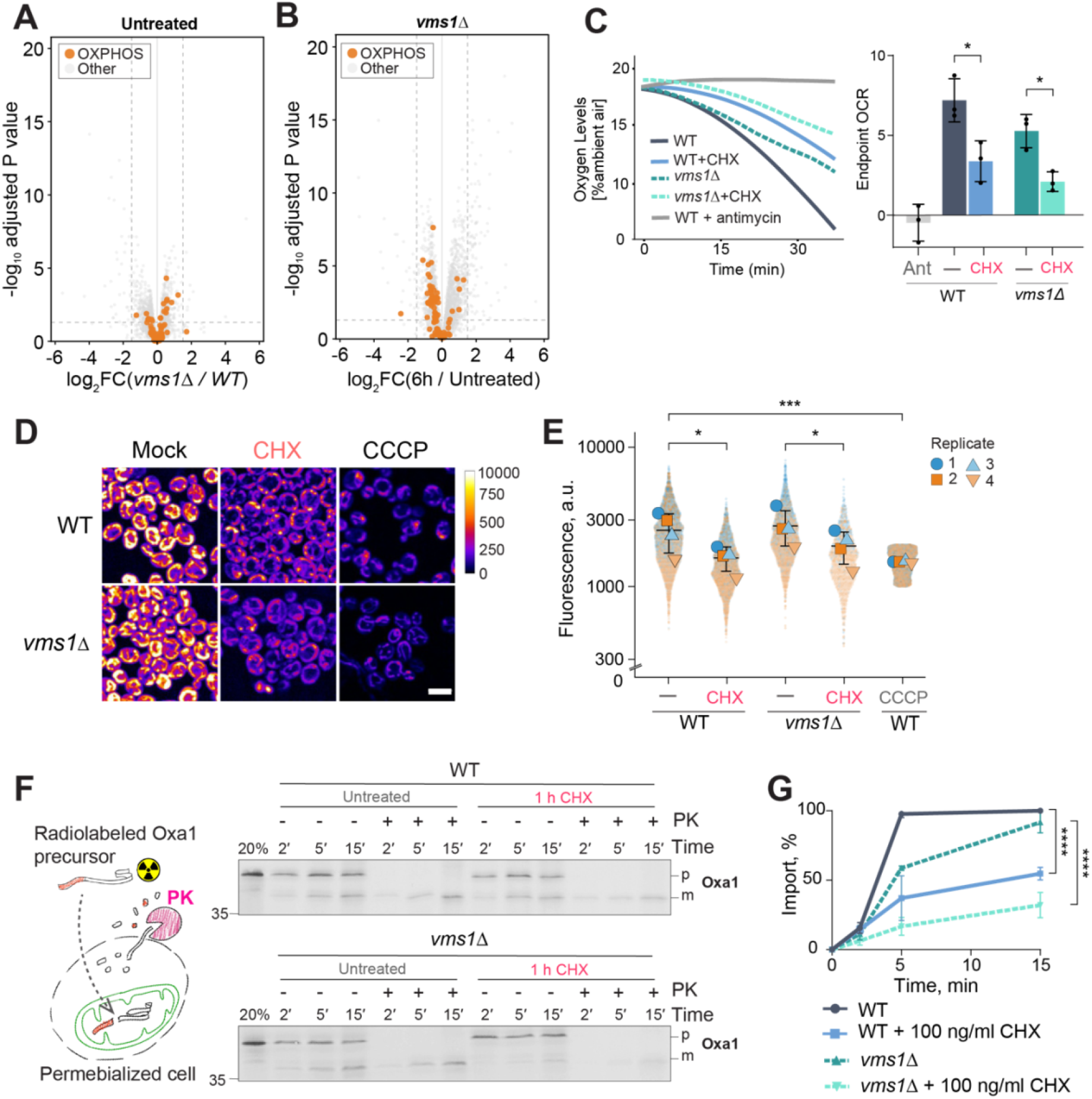
Cytosolic translation stress impairs mitochondrial respiration and import. **(A,B)** Volcano plots showing proteome changes in *vms1*Δ cells relative to untreated WT cells **(A)** and after 6 h of CHX treatment relative to untreated *vms1*Δ cells **(B)**. Log_2_ fold changes are plotted against −log_10_-adjusted P values. OXPHOS proteins are highlighted in orange **(C)** Oxygen consumption of WT and *vms1*Δ cells with or without 100 ng/ml CHX for 4 h. Antimycin-treated WT cells served as a respiration-deficient control. Oxygen consumption was measured in YPEtOH using FirePlate-O_2_ sensors and is shown as kinetics (right) and endpoint OCR values (left). Data represent three biological replicates. Statistical significance was determined by two-way ANOVA with Tukey’s multiple-comparisons test. **(D)** WT and *vms1*Δ cells were treated with 100 ng/ml CHX or 10 µM CCCP for 4 h and stained with 5 nM MitoTracker Red CMTXRos. Scale bar, 5 µm. **(E)** Quantification of single-cell fluorescence intensities from (D). Symbols indicate four independent biological replicates. Statistical significance was determined by two-way ANOVA with Tukey’s multiple-comparisons test. **(F)** Import of radiolabeled Oxa1 into semi-intact WT and *vms1*Δ cells untreated or pretreated with CHX for 1 h. Following import for 2, 5, or 15 min, samples were treated with or without proteinase K and analyzed by SDS-PAGE and autoradiography. An input corresponding to 20% of the radiolabeled Oxa1 was loaded. Precursor (p) and mature (m) forms are indicated. **(G)** Quantification of Oxa1 import, normalized to the maximal signal. *P < 0.05, **P < 0.01, ***P < 0.001, ****P < 0.0001.

While *VMS1* deletion did not have a strong effect on the protein composition of the OXPHOS complexes compared to WT cells, the examination of the protein abundance changes upon CHX treatment reveals a consistent decrease of protein abundances after 4 to 6 h of treatment for many complex subunits (Fig. 3B, Fig. S4A). We next aimed to test if this reduction is also accompanied by the decrease in respiratory capacity. For this, we analyzed oxygen consumption rate (OCR) of the WT and *vms1Δ* cells after 4 h treatment with 100 ng/ml CHX. We find that *vms1Δ* cells have lower OCR compared to WT and that CHX treatment reduces the OCR in both strains (Fig 3C). Next, we tested if OCR reduction was accompanied by the reduction of mitochondrial membrane potential. The uptake of membrane potential-sensitive dye was reduced to the same extent in WT and *vms1Δ* cells after 4 h incubation with 100 ng/ml CHX suggesting that *VMS1* deletion did not create a stronger phenotype compared to WT cells, but that cytosolic translation stress induces a strong membrane potential reduction in both strains (Fig. 3D,E). These results further emphasize that translation stress has a broad effect on mitochondrial function, and some aspects such as OCR may be stronger affected in *vms1Δ* strain. The reduced respiratory capacity however does not explain the hypersensitive CHX phenotype exhibited by the *VMS1* mutant on all growth conditions.

While respiration is only required for growth on non-fermentable carbon sources, mitochondrial protein import is essential for survival. To compare the mitochondrial import capacity between WT and *vms1Δ* cells, we used semi-intact cell (SIC) import assay that allows to measure protein translocation into mitochondria preserved within cellular context (Hansen et al., 2018). Already after one hour of treatment with 100 ng/ml CHX, the SIC capacity to import a radiolabeled precursor was significantly reduced in WT cells and almost completely abolished in *vms1Δ* mutants. Thus, we identify an essential cellular process that is strongly affected by *VMS1* deletion.

### Stress-response pathways helping to maintain mitochondrial function in VMS1 mutant

Artificial inhibition of mitochondrial protein import by overexpression of slowly imported proteins also abolishes cell growth and activates specific stress responses (Boos et al., 2019). We therefore decided to check if CHX-induced mitochondrial import reduction is also accompanied by the activation of similar stress pathways. Indeed, using quantitative PCR (qPCR) we detected the increase of mRNA levels of proteasome activator *RPN4* (Fig. 4A). On the protein level, core proteasome subunit abundance was increased already in untreated *vms1Δ* cells and was elevated in all CHX-treated *vms1Δ* samples (Fig. S5). *RPN4* expression can be activated either by proteasome dysfunction, or through the heat shock responsive transcription factor Hsf1. Indeed, we also found that *SSA2* mRNA abundance was increased, suggesting the potential involvement of the heat shock pathway (Fig. 4B). Protein-level analysis of heat shock protein chaperones revealed no clear changes upon CHX treatment and only minor differences between WT and *vms1*Δ cells, involving specific members of the Hsp70 and small HSP families, including Ssa1/2, Hsp42, and Hsp26 (Fig. S5). The modest increase in SSA2 mRNA, comparable to that observed under moderate heat stress at 37°C, suggests that CHX-induced translation stress elicits only a limited heat shock response. The accompanying induction of proteasome- and heat shock-related pathways may therefore not result directly from impaired mitochondrial protein import, but instead reflect cytosolic proteostatic stress caused by the accumulation of aberrant translation products. (Choe et al., 2016). A more specific mitochondrial import stress is signaled through the transcription factor Pdr3 that strongly activates *CIS1* mRNA expression (Weidberg and Amon, 2018; Yuan et al., 2026). We indeed found that CHX-induced translation stress leads to a small but significant increase of *CIS1* mRNA levels both in WT and *vms1Δ* cells that is much lower compared to the increase induced by overexpression of an artificial inhibitor of mitochondrial import, cytochrome b2 bipartite targeting signal fused to tightly folded protein DHFR (b2-DHFR) (Fig. 4C). Taken together, our findings show that cytosolic translation stress strongly impairs mitochondrial protein import and elicits a response that overlaps with canonical import stress, although *CIS1* induction remains substantially lower than in the b2-DHFR control (Boos et al., 2019).

**Figure 4.**
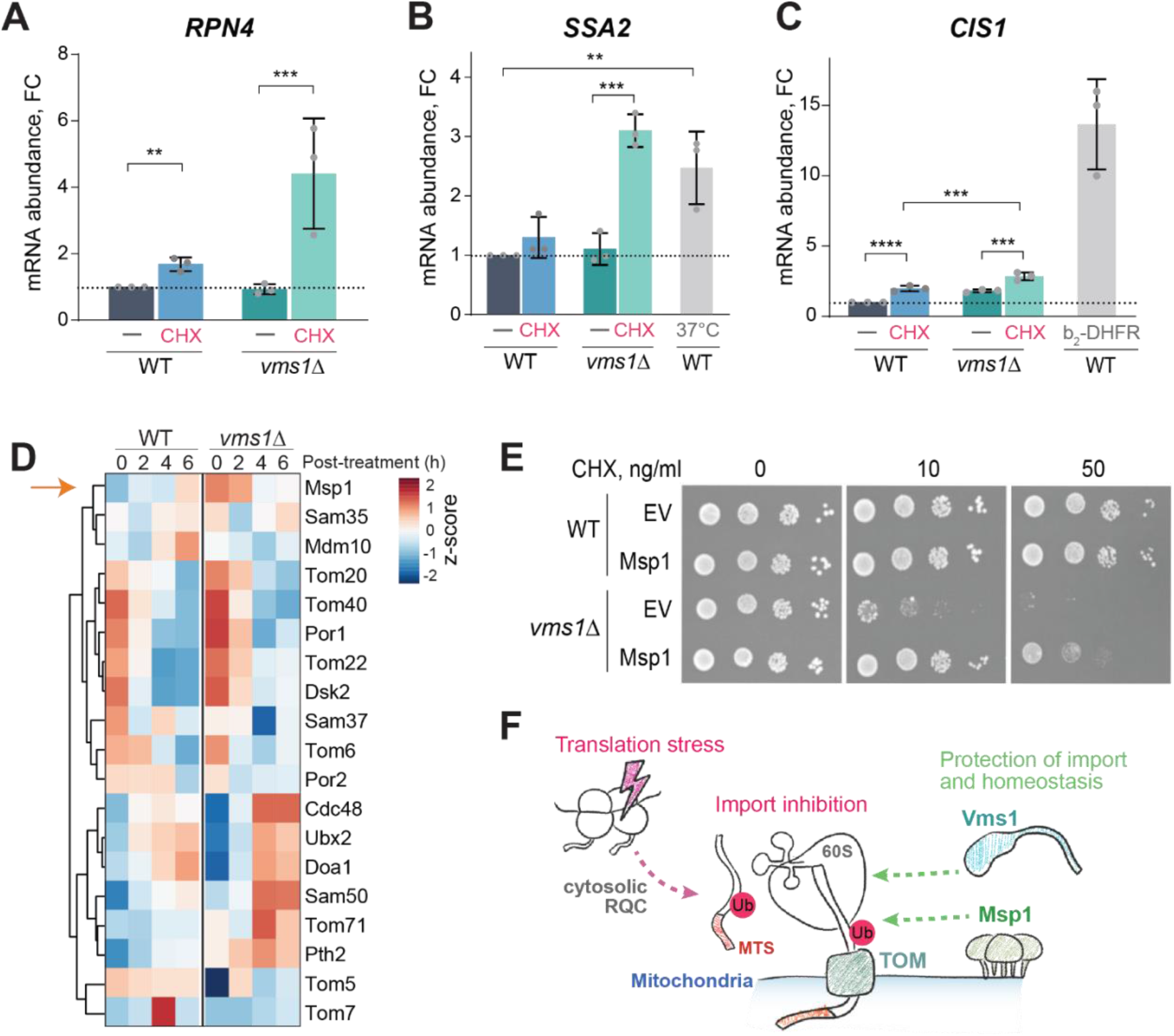
Loss of Vms1 activates cellular stress responses and alters the expression of mitochondrial quality control components. **(A–C)** Relative RPN4 (A), SSA2 (B), and CIS1 (C) mRNA levels in WT and *vms1Δ* cells treated with or without 100 ng/ml CHX for 4 h. Heat-shocked WT cells (37°C) and WT cells expressing b₂-DHFR served as positive controls for SSA2 and CIS1 induction, respectively. Values are shown relative to untreated WT cells. Points represent biological replicates. Statistical significance was determined by two-way ANOVA with Tukey’s multiple-comparisons test. **(D)** Heatmap of mitochondrial outer-membrane and quality-control proteins in WT and *vms1Δ* cells after 0–6 h of CHX treatment. Row-normalized z-scores and hierarchical clustering are shown. Msp1 is marked by an orange arrow. **(E)** Serial-dilution growth assay of WT and *vms1Δ* cells expressing Msp1 or carrying an empty vector (EV) on the indicated carbon sources and CHX concentrations. **(F)** Cytosolic translation stress can lead to the production of aberrant translation products as a result of cytosolic RQC pathway. Such ubiquitinated or 60S-bound nascent chains that contain a mitochondrial targeting signal (MTS) may block mitochondrial protein import by obstructing the translocase of the outer membrane (TOM) complex. Vms1 and Msp1 help to alleviate this blockage and rescue cell viability. *P < 0.05, **P < 0.01, ***P < 0.001, ****P < 0.0001.

To gain further insights into the effects of cytosolic translation stress on mitochondrial import quality control pathways, we revisited our proteomic data and examined the abundance dynamics of translocases and associated quality control factors that take part in mitoTAD and mitoCPR (Mårtensson et al., 2019; Schulte et al., 2023; Weidberg and Amon, 2018). The TOM complex subunits Tom40, Tom20, and Tom22 decreased in abundance after 4 h of CHX treatment in both WT and *vms1*Δ cells, mirroring the pattern observed for TCA-cycle enzymes and subunits of OXPHOS complexes II–V. (Fig S4A, Fig 4D). Several mitoTAD components, including Ubx2, Doa1, and Pth2, increased in both WT and *vms1*Δ cells, with a slightly stronger increase in *vms1*Δ. Msp1, a component of the mitoCPR pathway, was more abundant in *vms1*Δ at baseline, but this difference was no longer evident after 6 h of CHX treatment. The *MSP1* mRNA levels showed a tendency to increase in *vms1*Δ following CHX treatment, suggesting differences in the temporal regulation of *MSP1* transcript and protein abundance (Fig. S6A). Msp1 is an AAA-type ATPase that can clear stalled precursor proteins from the TOM complex and extract mislocalized tail-anchored proteins from the outer mitochondrial membrane (Chen et al., 2014; Kim et al., 2024). To test if Msp1 can alleviate stress, we overexpressed WT Msp1 or ATPase-defective mutant Msp1 E193Q in *vms1Δ* cells. Only WT Msp1 but not the mutant was able to improve the growth of *vms1Δ* strain on low CHX concentration showing that the extractase activity of Msp1 is required to overcome the deleterious effects of translation stress (Fig. 4E, Fig. S6B). We also found that overexpression of WT Msp1 alleviated the respiratory defect of CHX-treated *vms1Δ* cells, although the OCR remained lower than in untreated cells (Fig. S6C).

Our results suggest that aberrant cytosolic translation specifically causes mitochondrial protein import defect that is aggravated by *VMS1* deletion and can be rescued by the Msp1 extractase, revealing further connections between co-translational and post-translational protein quality control pathways (Fig. 4F).

## Discussion

In this work we investigated the toxic effects of translation stress and the role of Vms1 protein in promoting cell survival under this condition. To induce the stress, we used low concentrations of CHX that reversibly binds the E-site of the ribosome (Garreau de Loubresse et al., 2014). Unlike commonly used stalling reporters, such as non-stop proteins, CHX treatment can mimic the stresses that induce ribosome stalling at random positions, such as oxidative or chemical stress that damages mRNAs. Screening the yeast collection of non-essential gene mutants and the DAmP collection of hypomorphic alleles uncovered a unique CHX hypersensitivity yeast cells lacking peptidyl-tRNA hydrolase Vms1 (Fig. 1D). A similar phenotype has been reported previously for *vms1Δ* as part of the characterization of RQC mutants, where it was attributed to the general role of Vms1 in recycling 60S ribosomal subunits (Verma et al., 2018; Zurita Rendón et al., 2018). However, we did neither observe an increased accumulation of 60S subunits nor a decreased translation in CHX-treated *vms1Δ* cells compared to WT cells (Fig. 1E, Fig. S1E). This might be explained by the presence of alternative tRNA hydrolases such as Pth2 and Jlp2 that can recycle 60S (Bertram et al., 2025; Iyer et al., 2025). Supporting this notion, our global proteome analysis revealed upregulation of Pth2, Jlp2, and the ribosome splitting factor Dom34 in *vms1Δ* mutants (Fig. 2B), accompanied by broad upregulation of amino acid synthesis pathways (Fig. S3).

Our global proteome analyses further revealed a specific depletion of mitochondrial proteins in CHX-treated cells (Fig. 2). Consistent with this, mitochondrial protein import is significantly reduced in *vms1Δ* mutants (Fig. 3F). Notably, *vms1Δ* strain showed stronger import inhibition compared to WT. This functional defect may explain the cell death on all carbon sources (Fig. 1B) since mitochondrial protein import is an essential process. Furthermore, we show that overexpression of OM-localized AAA-ATPase Msp1 rescues CHX hypersensitivity of *vms1Δ* strain. This findings further link translation stress with mitochondrial import because the main functions of Msp1 are to remove mislocalized OM proteins and to clear blocked translocons (Chen et al., 2014; Kim et al., 2024; Wohlever et al., 2017). Overall, our data shows an unexpected and strong connection between cytosolic translation stress and mitochondrial fitness. We suggest that the decline of mitochondrial function is caused by reduced protein import into the organelle. We hypothesize that the Vms1 protein plays a special role in protecting mitochondrial import and ensuring cell survival under translation stress (Fig. 4F).

How can translation stress in the cytosol lead to the mitochondrial import defects and what is the role of Vms1? We suggest that there are three possible reasons which are not mutually exclusive: (1) the aggregation of CAT-tailed proteins in the matrix, (2) accumulation of TOM-bound 60S-nascent chain complexes, (3) accumulation of TOM-bound ubiquitinylated precursors. The specific role of Vms1 in protecting mitochondrial proteostasis from CAT-tailed proteins was previously explained by the ability of Vms1 to displace Rqc2 from 60S subunits and to release degradable nascent chains to the matrix (Izawa et al., 2017). The deletion of Vms1 leads to the aggregation of CAT-tailed proteins together with other matrix proteins. Matrix protein aggregation can lead to sequestration of mtHsp70 away from the TIM23 complex leading to reduced mitochondrial import (Banerjee et al., 2026). We cannot completely rule out the influence of CAT-tails, but our data does not fully agree with this explanation. Namely, the effect of CAT-tail accumulation in *vms1Δ* is enhanced by the deletion of Ltn1 and is observed only on respiratory media (Izawa et al., 2017), while the CHX-hypersensitivity of *vms1Δ* is alleviated by Ltn1 deletion and occurs on all carbon sources (Fig. 1B). Additional explanations for the observed mitochondrial import deficiency can be provided by the direct blockage of the TOM complex. For example, it was shown that overexpression of non-stop mitochondrial proteins in the strain with double deletion of Ski7 and Dom34 leads to mitochondrial import defects (Izawa et al., 2012). The defect may stem from direct blocking of the TOM complex by precursor-bound RNCs that cannot be disassembled. In case of Vms1 deletion the blockage may be mediated by 60S-nascent chain complexes that cannot be resolved. This possibility needs additional investigation to measure the association of 60S with TOM, elucidate why other tRNA-hydrolases cannot resolve it, and explain why Ltn1 deletion alleviates the phenotype. Finally, we suggest that the direct blockage of the TOM complex can be the consequence of nascent chain ubiquitination. Ubiquitin attached to the precursor proteins can block their import and occupy the TOM complexes. This can happen both with the nascent chains still attached to the 60S subunits, and to the nascent chains that are released but not degraded yet. This model can explain how Ltn1 deletion alleviates the CHX sensitivity of *vms1Δ* mutant by preventing translocon clogging with ubiquitin. Rqc2 deletion may be also alleviating the phenotype because this factor not only produces CAT-tails but also binds Ltn1 and enhances its stability and ubiquitin-ligase activity (Brandman et al., 2012; Lyumkis et al., 2014; Shao et al., 2015). Indeed, it was shown that the *vms1Δ* strain CHX-sensitivity depends on ubiquitin-ligase activity of Ltn1 but not on CAT-tailing activity of Rqc2 (Zurita Rendón et al., 2018). The ubiquitin-dependent blockage model also explains the strong genetic interaction of *VMS1* with *MSP1* (Fig. 4) and with mitoTAD component *UBX2* (Mårtensson et al., 2019). How does Vms1 prevent such toxic over-ubiquitinylation of mitochondrial precursor proteins by Ltn1, remains to be determined. Overall, our data emphasize a strong connection between cytosolic translation quality control and the mechanisms that safeguard mitochondrial proteostasis.

Cytosolic translation and mitochondrial proteostasis are tightly connected, both in normal and stress conditions. Protein synthesis in the cytosol and in the mitochondria must be coordinated to control the stoichiometry of molecular complexes encoded in both genomes (Couvillion et al., 2016). The retrograde stress signaling from mitochondria to cytosol is an active area of research. In mammalian cells mitochondrial dysfunction signal is transmitted through the integrated stress response pathway (ISR) by the activation of HRI kinase that phosphorylates translation initiation factor eIF2alpha (Fessler et al., 2020; Guo et al., 2020). In yeast, the mitochondrial import blockage triggers inhibition of translation and downregulation of mitochondrial proteome via HAP2/3/4 complex, but the mechanisms are less clear (Boos et al., 2019; Wrobel et al., 2015). At the same time, the accumulation of precursors in the cytosol activates CIS1 expression under the control of Pdr3 transcription factor (Weidberg and Amon, 2018). Pdr3 activation requires direct binding of Mge1 precursor mitochondrial targeting signal (Yuan et al., 2026). Low activation of *CIS1* expression during CHX-induced mitochondrial import defect that we observe can be explained by decreased translation and increased proteasome abundance that can reduce the amount of available Mge1 precursor in the nucleus.

Our discovery that cytosolic translation stress can cause mitochondrial dysfunction expands the known retrograde cytosol-mitochondria connection in the other direction. It remains to be determined whether the observed decline of mitochondrial protein levels results from reduced protein import, and adaptive response, or both and whether a similar occurs in mammalian cells. The main response induced by translation stress is detected by ribosome collisions. In mammalian cells, collided ribosomes trigger the ISR via GCN2 and the ribotoxic stress response via ZAKalpha (Wu et al., 2020). While the ISR reduces collisions and rescues stalled ribosomes, ZAK signaling promote either survival or apoptosis (Yan and Zaher, 2021; Nanjaraj Urs et al., 2024; Sinha et al., 2024). In yeast, the GCN2-dependent ISR triggered by ribosome stalling is effectively counteracted by the activity of RQC and may not lead to significant adaptations when the collision rate is low (Nanjaraj Urs et al., 2024). In the future it would be interesting to find out how mitochondrial function is affected by these signaling pathways and how they interact with mitochondrial stress signaling in both yeast and mammalian cells.

Ribosome stalling by low concentrations of inhibitors is a useful model that allows to investigate translation stress separately from other stressors, such as oxidative agents, or starvation, that may also cause ribosome collisions. However, for free-living unicellular eukaryotes, such as yeast, the presence of antibiotics is a physiologically relevant situation since these compounds are produced by bacteria and are present in the environment. Investigation of antibiotic effects is also important for the development of antifungal drugs. Here we show that *VMS1* deletion renders yeast highly sensitive to translation inhibitors and that the effect can be counteracted by mitochondrial quality control protein Msp1. We find that the cells exposed to CHX might die from the combination of translation and mitochondrial import stress. These findings raise the possibility that simultaneous disruption of cytosolic translation and mitochondrial quality control could provide an effective strategy for antifungal combination therapies (Li et al., 2025b; Vallières et al., 2018).

Development of combination antifungal therapies is particularly attractive because fungi have efficient defense mechanisms such as pleiotropic drug resistance (PDR) pathway (Buechel and Pinkett, 2020). PDR is activated by transcription factors Pdr1 and Pdr3 that induce the expression of efflux pumps that remove the dangerous compounds. Pdr1 may directly sense xenobiotics such as cycloheximide (Buechel and Pinkett, 2020; Gale et al., 2023). Pdr3 is also responsible for the activation of mitoCPR by direct binding of unimported Mge1 precursor (Yuan et al., 2026). The evolutionary logic underlying the integration of mitochondrial stress responses with the PDR may lie in the fact that many xenobiotics directly target mitochondrial function (Li et al., 2025b). Our finding that cytosolic translation stress causes mitochondrial import defects suggests an additional explanation: compounds that primarily target cytosolic translation can also indirectly damage the mitochondria, creating a selective advantage for coordinating xenobiotic defense with mitochondrial stress responses. Overall, our study reveals previously unrecognized link between cytosolic translation stress and mitochondrial proteostasis mechanisms suggesting potential approaches for the developing of antifungal therapies.

## Acknowledgements

We are grateful to Christian Koch, Katja Hansen, and Lorenz Spänle for the help with experiments and to Sabine Knaus, Vera Nehr, Regina Jerhof and Cornelia Parent for technical support. We also would like to thank Emma Fenech for critical reading and editing of the manuscript and all the members of Bykov and Herrmann group for useful discussions. We thank Johannes Herrmann, Jared Rutter and Shunsuke Matsumoto for generously sharing plasmids, and Maya Schuldiner for sharing the yeast mutant collection and the WT strain. Research in the Bykov group is funded by BioComp initiative of the Land Rhineland-Palatinate, the Deutsche Forschungsgemeinschaft (DFG, German Research Foundation) – SPP2453 project number 541626282, and by the European Union (ERC StG, 3DTOP, 101165504). Research in the den Brave group is funded by priority program SPP 2453 BR 6283/6-1 project ID 541596792 and BR 6283/5-1 project ID 529716110. Research in Breker group is funded from the European Research Council Starting Grant (ERC-StG), project Chloro-Import (grant no. 101117931). Views and opinions expressed are, however, those of the authors only and do not necessarily reflect those of the European Union or the European Research Council. Neither the European Union nor the granting authority can be held responsible for them.

## Data Availability

Proteomics dataset is available via ProteomeXchange with identifier PXD081279. Raw sequencing data is available from the NCBI SRA database, identifier PRJNA1493039. All other data is included in the submission.

## Reviewer access details for ProteomeXchange

Log in to the PRIDE website using the following details:

## Project accession

PXD081279

Token

Mzt9q6KaIcMt

## Materials and Methods

### Strains and growth conditions

The yeast strains and plasmids used in this study are described in detail in Supplementary Table S3 and S4, respectively. Unless specified, all strains were derived from S288C MATα {*his3Δ1 leu2Δ0 met15Δ0 ura3Δ0 lys2+/lys+ can1Δ::GAL1pr-SceI::STE2pr-SpHIS5 lyp1Δ::STE3pr-LEU2*} [phi+]. The knock out strains were generated with a standard PCR-based gene editing followed by Li-Ac/ssDNA/PEG-based transformation (Gietz and Woods, 2006). The resistance genes were amplified from the standard plasmid sets (Longtine et al., 1998; Janke et al., 2004) using the primers listed in Table S5. Primer sequences were designed using Primers-4-Yeast (Yofe and Schuldiner, 2014). All knock outs were verified by PCR. The WT and *vms1*Δ strains (YYB038 and YYB055) were additionally verified by whole-genomic sequencing (see next section and Fig. S7).

The strains were grown at 30°C either in yeast complete medium (YP) containing 1% (w/v) yeast extract, 2% (w/v) peptone and 2% (w/v) of the respective carbon source or in minimal synthetic respiratory medium containing 0.67% (w/v) yeast nitrogen base, optimized OMM amino acid mix (Hanscho et al., 2012) and 2% (w/v) of the respective carbon source.

### Whole-genome sequencing and variant analysis

To make sure that there are no additional mutations that cause *vms1Δ* strain phenotype we verified the genomic DNA sequence of WT strain (YYB038) and *vms1Δ* strain (YYB055) used in this study using whole-genomic sequencing. Genomic DNA was purified from 10 ml overnight culture using 10 min lysis with glass beads in a vortex followed by phenol-chloroform extraction. The DNA samples were submitted to Azenta Lifesciences for library preparation and sequencing with Illumina NovaSeq XPlus platform at 2×150bp configuration. Raw reads were trimmed with fastp v1.0.1 (Chen, 2025) to remove poly-G tails, adapter sequences, and low-quality bases, retaining reads longer than 35 bp. Trimmed reads were mapped to the *S. cerevisiae* strain S288C reference genome (release R64-5-1; (Engel et al., 2025)) with BWA-MEM v0.7.19 (Li, 2013), and duplicate reads were removed with Picard v.3.4.0 (https://broadinstitute.github.io/picard/). SNPs and short indels were called with FreeBayes v1.3.10 (Garrison and Marth, 2012), requiring a minimum mapping quality of 30 and base quality of 20. Calls were decomposed and left-normalized against the reference with bcftools v1.23 (Danecek et al., 2021) and retained only if they had QUAL ≥ 30, total depth ≥ 10, alternate-allele support on both strands (SAF, SAR > 0) and both read ends (RPR, RPL > 1), and a quality-to-alternate-observation ratio ≥ 10; a variant was kept if, in at least one sample, the alternate allele was supported by ≥ 3 observations at ≥ 10× depth with an alternate-allele fraction ≥ 0.8. BY4741-background variants were obtained from the SGD strain archive(http://sgdarchive.yeastgenome.org/sequence/strains/BY4741/BY4741_Stanford_2014_JRIS00000000/), which provides call sets for BY4741 generated against the R64-1-1/sacCer3 reference (GenBank JRIS00000000). After removing these variants, no missense or nonsense variants distinguished the wild type from the *vms1Δ* strain; the remaining six missense variants were common to both.

Large indels were called with Delly v1.5.0 (Rausch et al., 2012) and verified manually in IGV (Robinson et al., 2011). All identified indels, except VMS1 deletion, were common in both strains. The raw reads were deposited to the NCBI SRA database, PRJNA1493039. The comparison of the WT and *vms1Δ* strain did not reveal any de novo mutations except the deletion of the *VMS1* gene (Fig. S7).

### Growth assays and viability tests

For spot analysis, the respective yeast strains were grown in YPGal media. 1 OD unit (amount of culture equivalent to 1 ml of culture with OD_600_=1.0) of cells in logarithmic growth phase (OD_600_=0.5-0.7) were harvested at the exponential phase. The cells were washed in sterile water and 3 µl of ten-fold serial dilutions were spotted on the respective media followed by incubation at 30°C. Scans were taken after 2 days (for fermentative media without CHX) or 3 days (respiratory media, or any CHX-containing media).

Growth curves were recorded in a 96 well plate, using the automated *SPECTROstar^Nano^* by *BMG Labtech*. Before starting the growth curves, cells were grown to logarithmic phase in YPGal (OD_600_=0.5-0.7) and diluted to OD_600_=0.2. The OD_600_ was measured every 10 min for 48 h at 30°C. The average of technical triplicates was calculated and plotted in Prism GraphPad.

For survival assays cells were grown in liquid YPGal media and optionally treated with cycloheximide for 2, 4, or 6 h. The cells were diluted to OD_600_=0.001 and 100 µl were plated onto YPGalactose plates and incubated for 2 days. The plates were photographed using a smartphone and the colonies were counted using Promega colony counter app (https://apps.apple.com/us/app/promega-colony-counter/id620431249).

### Mutant collections screen

A genetic screen was performed using the *Saccharomyces cerevisiae* knockout (KO) collection (Giaever et al., 2002) and the decreased abundance by mRNA perturbation (DAmP) collection (Breslow et al., 2008). Using a ROTOR+ pinning robot (Singer Instruments), strains were initially arrayed on solid glucose-containing medium in 1536 format and incubated for 2 days. Colonies were then replica-pinned onto solid YP medium supplemented with either glucose or glycerol as the carbon source. For each growth condition, parallel plates containing 100 ng/mL cycloheximide (CHX) were prepared. Plates containing glucose were incubated for 2 days, whereas plates containing glycerol were incubated for 3 days. After incubation, the plates were scanned and colony sizes were quantified using the SGAtools web platform (http://sgatools.ccbr.utoronto.ca/) (Wagih et al., 2013). CHX-dependent growth phenotypes under respiratory conditions were evaluated by comparing colony sizes on YPGlycerol medium in the presence and absence of CHX (Table S1).

### Oxygen consumption measurements

To measure cellular oxygen consumption over time, yeast cells were grown in YPGal medium. At an OD_600_ of 0.7, cells were harvested by centrifugation, washed with ddH_2_O, and resuspended in 125 μL YP-EtOH. Oxygen levels were measured using a FirePlate-O_2_ optical sensor system (PyroScience). Sensors were calibrated in ambient air according to the manufacturer’s instructions, and oxygen values are reported as percentage air saturation, with ambient air corresponding to approximately 21% O_2_.

After calibration, the cell suspension was transferred to the sensor plate, and an additional 190 μL YP-EtOH was added. Oxygen levels were recorded for 20 min. As a control, antimycin was added.

### Western blotting

For analysis of whole cell extracts by western blotting, 3 OD units were harvested by centrifugation (5 min, 5000 g, room temperature). Pellets were incubated with 0.2 M NaOH for 5 min at room temperature followed by centrifugation (5 min, 5000 g, room temperature). The pellet was resuspended in 150 µl HU sample buffer (5 % SDS (w/v), 8 M Urea, 1 mM EDTA, 200 mM Tris-HCl pH 6.8, 0.025 (w/v) mM bromphenol blue). Samples were analyzed by western blotting. The antibody against Tom22 was raised in rabbit (TB557). Ubiquitin was detected with monoclonal mouse anti Ubiquitin (Clone P4D1, Santa Cruz, sc-8017). Fluorescent secondary antibodies were from LiCor (goat anti-rabbit IgG, IRDye 800CW Cat. #: 926-32211 and goat anti-mouse IgG, IRDye 800CW, Cat. #: 926-32210). Images were taken on an LiCor Odyssey CLx Imager.

### Polysome fractionation on sucrose gradients

Yeast cultures were inoculated into 1 L of YPGal at an OD_600_ of 0.01 and grown to an OD_600_ of approximately 1.0. In addition, yeast cultures were treated during growth with 100 ng/ml cycloheximide (CHX) for 2, 4, or 6 h. Before harvesting, CHX was added to a final concentration of 0.05 mg/ml to stabilize the polysomes. The cells were harvested by centrifugation 5 min later. Cell pellets were washed with polysome extraction buffer containing 20 mM Tris-HCl pH 7.4, 50 mM KCl, 10 mM MgCl_2_, 1 mM dithiothreitol (DTT), 100 μg/ml cycloheximide, 1× RNase Inhibitor Murine (NEB), and 1× yeast protease inhibitor cocktail (Roche) diluted according to manufacturer instructions. Pellets were resuspended in 700 μl extraction buffer, and an equal volume of 1 mm glass beads was added. The suspensions were transferred to screw-cap tubes and lysed using a FastPrep-24 5G homogenizer (MP Biomedicals, Heidelberg, Germany) for three cycles of 30 s at 8.0 m/s, with 120 s breaks between cycles. Lysates were centrifuged at 3000 × g for 5 min at 4°C. The supernatant was subsequently centrifuged at 11300 × g for 2 min at 4°C, followed by an additional centrifugation step at 11300 × g for 10 min at 4°C. Clarified extracts corresponding to 300 μg of RNA were layered onto linear 10–50% sucrose gradients and centrifuged at 260,000 × g for 90 min in an SW41Ti swinging-bucket rotor (Beckman). Gradients were prepared and fractionated using a Biocomp Gradient station and Piston Gradient Fractionator. Absorbance at 260 nm (A_260_) was recorded.

### Analysis of mRNA levels by qRT-PCR

For total RNA extraction yeast strains were cultivated in synthetic media to mid-log phase. 4 OD units of cells were harvested for RNA extraction using the RNeasy Mini Kit (Qiagen) in conjunction with the RNase-Free DNase Set (Qiagen) according to the manufacturer’s instructions. Yield and purity of the obtained RNA was determined with a Spectrophotometer/Fluorometer DS-11 FX+ (DeNovix). 500 ng RNA were reverse transcribed into cDNA using the qScript cDNA Synthesis Kit (Quanta Bioscniences) according to the manufacturer’s instructions. To measure relative mRNA levels, the iTaq Universal SYBR Green Supermix (BioRad) was used with 2 µl of a 1:10 dilution of cDNA sample. For assessment of rRNA in isolated mitochondria the Luna Universal Probe One-Step RT-qPCR Kit (NEB) was used with 2 µl of RNA sample. Measurements were performed in technical triplicates with the CFX96 Touch Real-Time PCR Detection System (BioRad). Calculations of the relative mRNA expressions were conducted following the 2^-ΔΔCt^ method (Livak and Schmittgen, 2001). For normalization, the housekeeping gene *ACT1* was used due to its stability. See Table S5 for primer sequences.

### Isolation of semi-intact cells

The SIC preparation was performed essentially as described (Hansen et al., 2018). Precultures grew in SGal or YPGal at 30°C and were harvested (700 × g, 7 min, RT) in the exponential phase. The cell pellet was resuspended in 25 ml SP1 buffer (10 mM DTT, 100 mM Tris pH unadjusted) and incubated for 10 min at 30°C shaking. After centrifugation (1000 × g, 5 min, RT) the pellet was resuspended in 6 ml SP2 buffer (0.6 M sorbitol, 1 × YP, 0.2% glucose, 50 mM KPi pH 7.4, 3 mg/g wet weight zymolyase) and incubated at 30°C for 30-60 min. Spheroplasts were collected and resuspended in 40 ml of SP3 buffer (1x YP, 1% glucose, 0.7 M sorbitol) and incubated for 20 min at 30°C shaking. After centrifugation (1000 × g, 5 min, 4°C) spheroplasts were washed two times with 20 ml of ice cold permeabilization buffer (20 mM Hepes pH 6.8, 150 mM KOAc, 2 mM Mg-Acetate, 0.4 M sorbitol). The pellet was resuspended in 1 ml permeabilization buffer containing 0.5 mM EGTA and 100 µl aliquots were slowly frozen over liquid nitrogen for 30 min.

### Import into semi-intact Cells

Semi-intact cells were thawed on ice and the OD_600_ was measured in 1.2 M Sorbitol. Semi-intact cells in the amount of 0.2 units were used per reaction. Semi-intact cells were added to a mixture of B88 buffer, 2 mM ATP, 2 mM NADH, 5 mM creatine phosphate and 100 µg/ml creatine phosphatase. Radiolabeled Oxa1 precursor was synthesized in vitro using TnT® Quick Coupled Transcription/Translation System kit (Promega) with the addition of ^35^S-methionine (1 μl of a 11 μCi solution). Radiolabeled lysate was added to SICs, and the mixture incubated 10 min on ice to allow the cells to take up the lysate. Afterwards the suspensions were incubated at 30°C. For kinetics samples were taken after 5 and 20 min. The import reaction was stopped at 1:10 dilution in ice cold B88 buffer containing 2 mM CCCP and treated with or without 100 µg/ml proteinase K for 30 min on ice. Protein digestion was stopped by the addition of 2 mM PMSF. Semi-intact cells were centrifuged (4000 × g, 5 min, 4°C), washed again with B88 and 2 mM PMSF and centrifuged for 10 min at 16000 × g and 4°C. Finally, the pellet was resuspended by reducing sample buffer and resolved via SDS-PAGE.

### Radioactive in vivo labelling of cytosolic translation products

Cells were grown in galactose medium lacking methionine to exponential phase. Aliquots of 2 OD units of cells were harvested and incubated in no amino acid containing media for 15 minutes. ^35^S-methionine (1 μl of a 11 μCi solution) was added to the cell suspension and newly synthesized proteins were labeled for 5 min. Incorporation of radioactive methionine was quenched by the addition of 8 mM cold methionine. Cells were lysed with 0.3 M NaOH containing 1% β-mercaptoethanol and 3 mM PMSF. Proteins were precipitated with 12% trichloroacetic acid and analyzed by SDS–PAGE and autoradiography.

### Quantitative whole-proteome analysis

Yeast cells were grown in YPGal at 30°C using four biological replicates per condition (n = 4). For each sample, 20 OD_600_ units were harvested by centrifugation at 5000 × g for 5 min, washed with ice-cold water, snap-frozen in liquid nitrogen, and stored at −80°C. Cells were lysed in a buffer containing 6 M guanidinium chloride, 10 mM TCEP-HCl, 40 mM 2-chloroacetamide (CAA; Sigma-Aldrich), and 100 mM Tris-HCl (pH 8.5). Mechanical disruption was performed with glass beads using a FastPrep-24 5G homogenizer (MP Biomedicals) for three 30-s cycles at 8.0 m/s, with 120-s breaks between cycles. Lysates were subsequently heated at 96°C for 10 min and clarified by centrifugation at 16,000 × g for 5 min at 4°C. Protein concentrations were determined using the Pierce BCA Protein Assay Kit (Thermo Scientific).

For peptide digestion, 25 µg of protein per sample was diluted with nine volumes of digestion buffer containing 10% acetonitrile and 25 mM Tris-HCl (pH 8.5). Trypsin (Promega, #V5111) and LysC (Wako, #125-05061) were each added at an enzyme-to-protein ratio of 1:50 (w/w), and samples were digested overnight at 37°C. The resulting peptides were acidified to pH < 2 with trifluoroacetic acid (TFA) and desalted using StageTips packed with three layers of SDB-RPS 3M disks. StageTips were activated with acetonitrile, equilibrated with 30% methanol and 1% TFA, and washed with 0.2% TFA. After sample loading, the peptides were washed again with 0.2% TFA and eluted with 80% acetonitrile and 5% ammonia. Eluates were dried by vacuum centrifugation and resuspended in buffer A containing 0.1% formic acid supplemented with 10% buffer A* containing 2% acetonitrile and 0.1% TFA.

Peptides were separated on in-house-packed 50-cm analytical columns with a 75-µm inner diameter containing ReproSil-Pur 120 C18-AQ resin with a particle size of 1.9 µm (Dr. Maisch). Liquid chromatography was performed using an EASY-nLC 1200 system coupled to a Q Exactive HF mass spectrometer (both Thermo Scientific). Peptides were eluted using a 3-h gradient from 2% to 95% buffer B, consisting of 80% acetonitrile and 0.1% formic acid, with 0.1% formic acid used as buffer A. Mass-spectrometric data were acquired using a Top15 method. Raw data were processed using MaxQuant v2.6.4.0. All additional acquisition and data-processing parameters are provided with the dataset deposited in the PRIDE repository.

Protein-group data generated by MaxQuant were processed in R v4.5.1 (R Core Team, 2020). Contaminants, reverse hits, proteins identified only by site, and proteins detected in fewer than three of the four biological replicates were excluded. After filtering, 3,715 robustly identified protein groups remained. Label-free quantification (LFQ) intensities were log₂-transformed, subjected to variance-stabilizing normalization, and corrected for batch effects using limma. Missing values were imputed only when no valid LFQ intensity was available for any replicate of a condition. In these cases, four values were sampled from a normal distribution using a fixed random seed of 7129456. For each dataset, the mean of the distribution was set to the first percentile of the LFQ-intensity distribution. Its standard deviation was derived from the median of the within-sample LFQ-intensity standard deviations calculated across the quadruplicate measurements. Differential protein abundance was assessed using limma for the indicated pairwise comparisons, and P values were adjusted for multiple testing using the Benjamini–Hochberg procedure. Principal component analysis was performed on the processed LFQ intensities using singular value decomposition. Mitochondrial sublocalization was assigned using a merged and manually curated list derived from published datasets by Vögtle and colleagues.

Gene Ontology enrichment analysis was performed in R using a custom script based on the enrichment strategy implemented in ShinyGO (Ge et al., 2020). Yeast systematic ORF identifiers were mapped to Entrez Gene identifiers using the org.Sc.sgd.db and AnnotationDbi packages. All successfully mapped proteins detected in the dataset were used as the background gene universe. For each comparison, significantly altered proteins were selected based on adjusted q values and abundance differences and subsequently divided into upregulated and downregulated protein sets. Enrichment analyses were performed separately for Biological Process and Cellular Component terms using a hypergeometric over-representation test. Only GO terms containing between 5 and 500 genes from the background universe were considered. P values were adjusted using the Benjamini–Hochberg method, and terms with a false-discovery rate of ≤ 0.05 were considered significantly enriched. Redundancy among enriched GO terms was reduced on the basis of semantic similarity using rrvgo with a similarity threshold of 0.7. Enriched terms were visualized as lollipop plots using ggplot2, displaying −log_10_(FDR), fold enrichment, and the number of overlapping genes.

### Fluorescence microscopy

Cells were grown in YPGal medium to logarithmic phase and treated with 100 ng/ml CHX for 4 h, or with 10 µM CCCP for 4 h. Cells were then transferred onto Concanavalin A (ConA)-coated glass-bottom plates and stained with 5 nM MitoTracker™ Red CMXRos (Invitrogen) for 10 min at room temperature. Samples were protected from light whenever possible to minimize photobleaching. Confocal fluorescence microscopy was performed using an SpinSR automated spinning disk microscope (Olympus/Evident) equipped with Yokagawa W1 scanning unit, multi-bandpass dichroic mirrorm and Orca Fusion sCMOS camera (Hammamatsu) under the control of ScanR Acquisition software. The images were acquired with a 20× air objective and a 3.2× magnification changer. MitoTracker Red was excited at 561 nm, and fluorescence emission was collected using 615/40 emission filter. Image visualization was performed using ImageJ/Fiji. The measurements of single-cell fluorescence intensity were performed using ScanR Analsysis software. Statistical analyses were performed in RStudio.

## Supplementary materials

**Table S1.** The comparison of growth rates of whole-genomic mutant collections under translation stress and normal growth conditions.

**Table S2.** Whole-cell proteome dynamics of WT and vms1Δ under translation stress.

**Table S3**. List of yeast strains used in this study.

**Table S4.** List of plasmids used in this study.

**Table S5**. List of primers used for yeast transformations.

---

**Figure S1.**
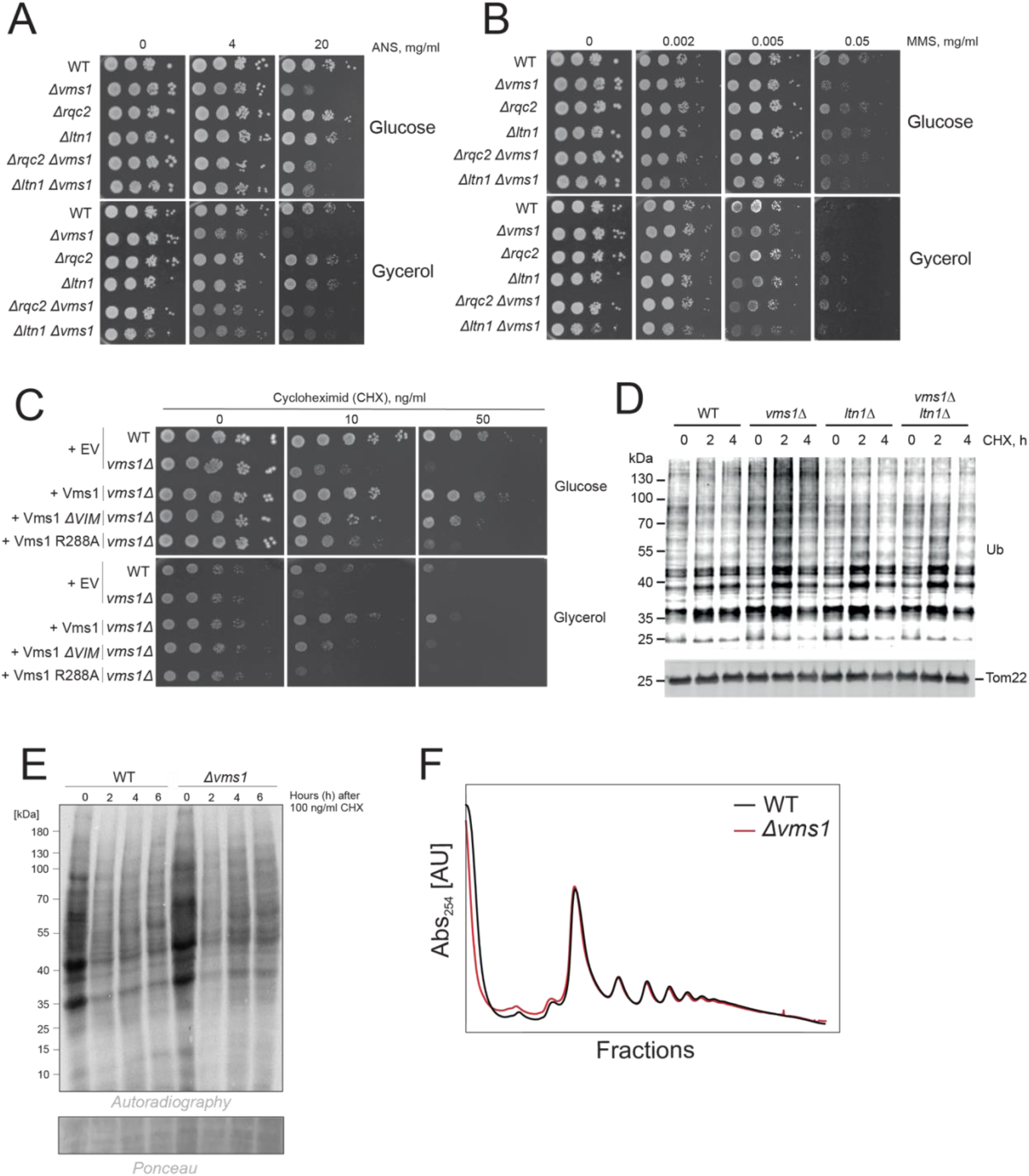
Vms1 modulates cellular stress tolerance and cycloheximide sensitivity. **(A, B)** Serial-dilution growth assays of the indicated RQC mutants on YPD and YPGly containing anisomycin (ANS) (A) or methyl methanesulfonate (MMS) **(B).** Plates were incubated at 30°C. **(C)** Serial-dilution growth assay of WT and *vms1Δ* cells carrying an empty vector (EV), or *vms1Δ* cells expressing Vms1, Vms1Δ*VIM*, or Vms1 R288A, on the indicated carbon sources and CHX concentrations. Plates were incubated at 30°C. **(D)** Western blot analysis of ubiquitinated proteins and Tom22 in the indicated strains after treatment with 100 ng/ml CHX for 2–6 h; 0 h represents the untreated control. **(E)** Cytosolic protein synthesis in WT and *vms1Δ* cells after CHX treatment, assessed by metabolic labeling with [³⁵S]methionine. Newly synthesized proteins were visualized by autoradiography, and Ponceau staining served as a loading control. **(F)** Polysome profiles of WT and *vms1Δ* cells grown in YPGal. Lysates were separated on 10–50% sucrose gradients, and RNA absorbance was recorded during fractionation.

**Figure S2.**
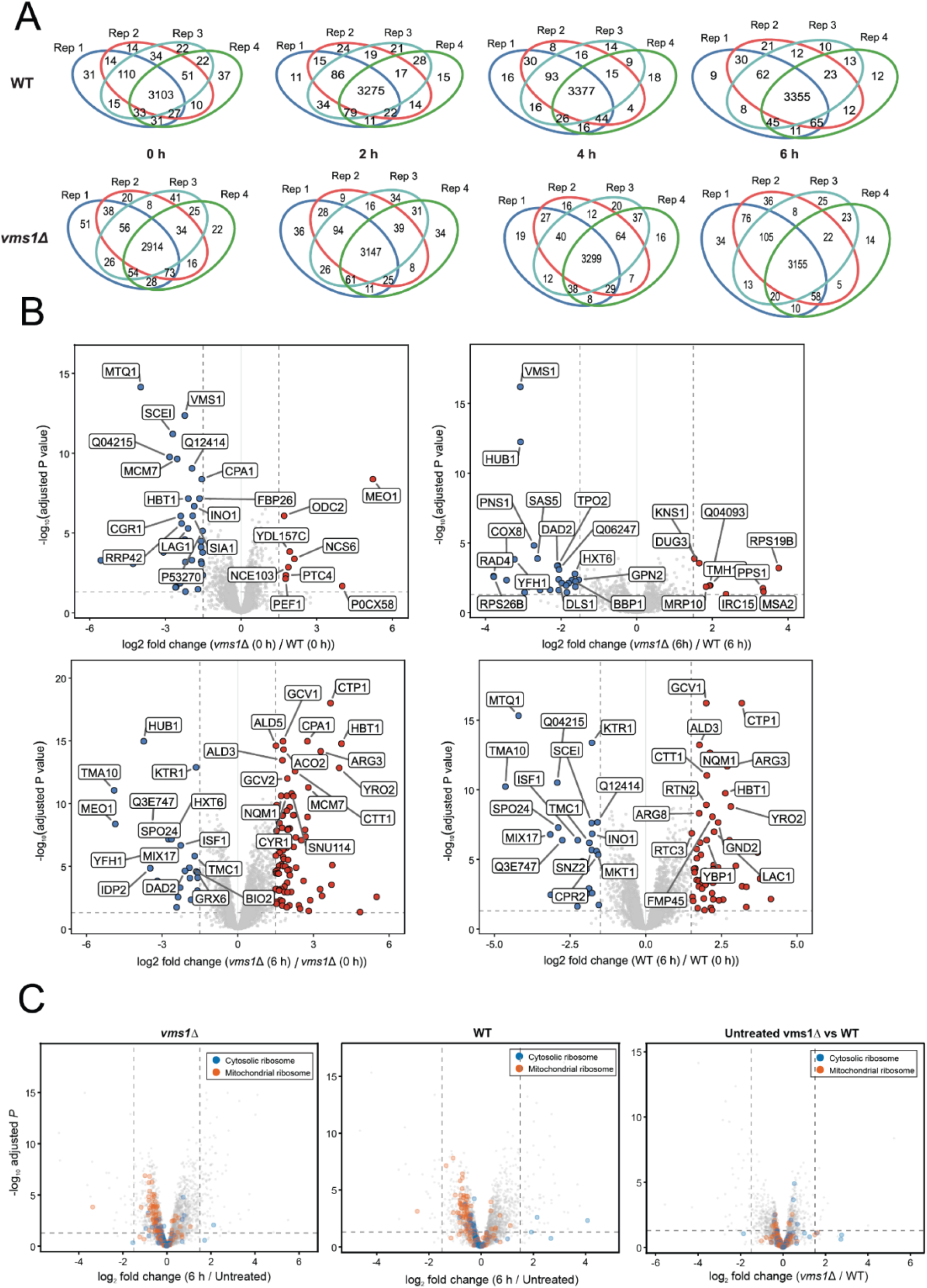
Time-resolved proteomic changes reveal an altered translational response in *vms1Δ* cells. **(A)** Venn diagrams showing proteins detected across four biological replicates of WT and *vms1*Δ cells after 0–6 h of CHX treatment. Numbers indicate proteins unique to or shared among replicates. **(B)** Volcano plot of proteome differences between untreated *vms1*Δ and WT cells. Significantly decreased and increased proteins are shown in blue and red, respectively, and selected proteins are labeled. **(C)** Volcano plots showing proteome changes after 6 h of CHX treatment relative to the respective untreated condition in *vms1*Δ (left) and WT cells (middle), and differences between untreated *vms1*Δ and WT cells (right). Cytosolic ribosomal proteins, and mitoribosomal proteins are highlighted in blue and orange, respectively. Log_2_ fold changes are plotted against −log_10_-adjusted P values.

**Figure S3.**
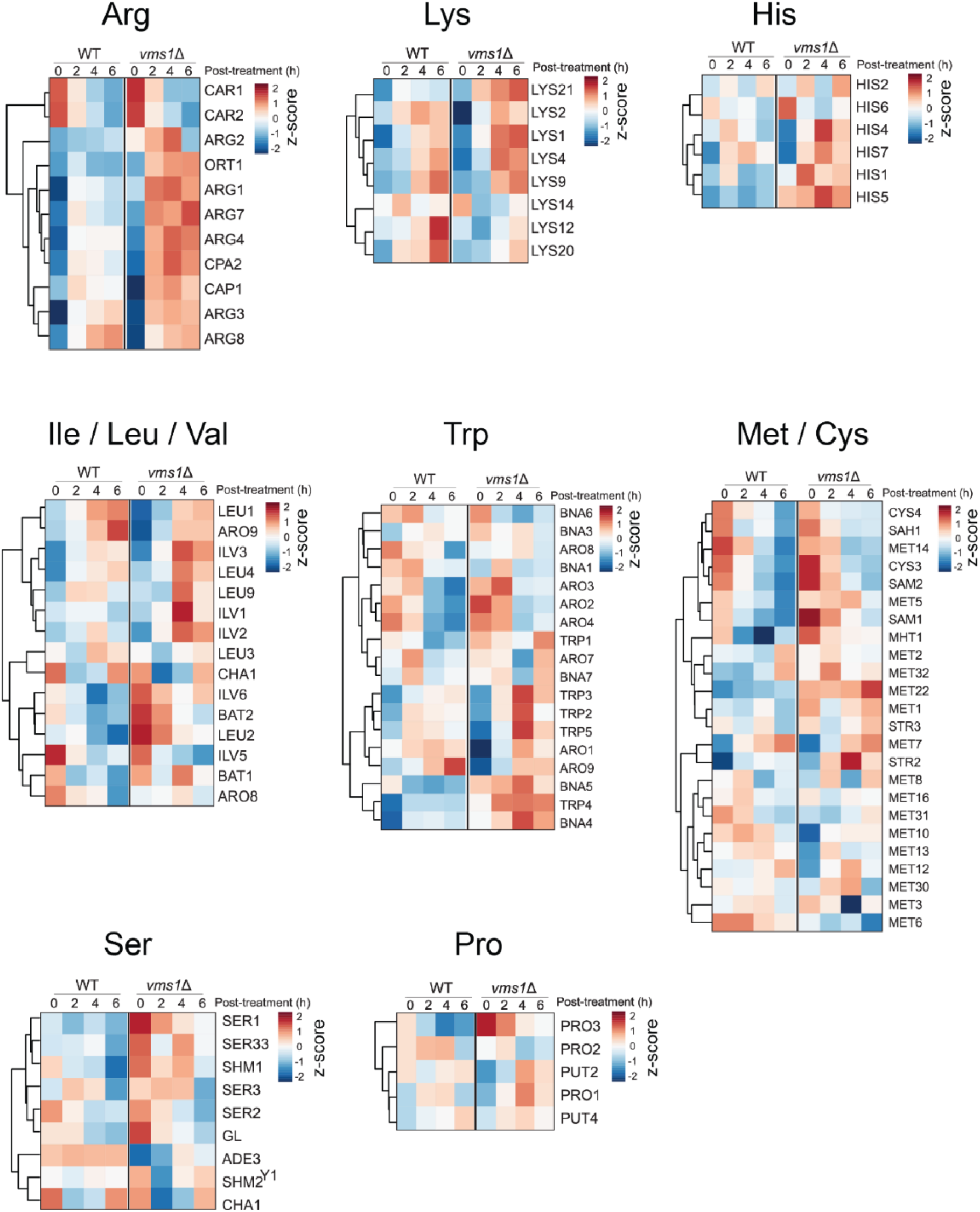
Loss of Vms1 alters the transcriptional response of amino acid biosynthesis and metabolism pathways. (A-I) Heatmaps showing the relative abundance of proteins associated with the indicated amino acid biosynthesis and metabolism pathways in WT and *vms1Δ* cells at 0, 2, 4, and 6 h after treatment. Values are row-normalized and displayed as *z*-scores. Proteins were hierarchically clustered according to their temporal expression profiles.

**Figure S4.**
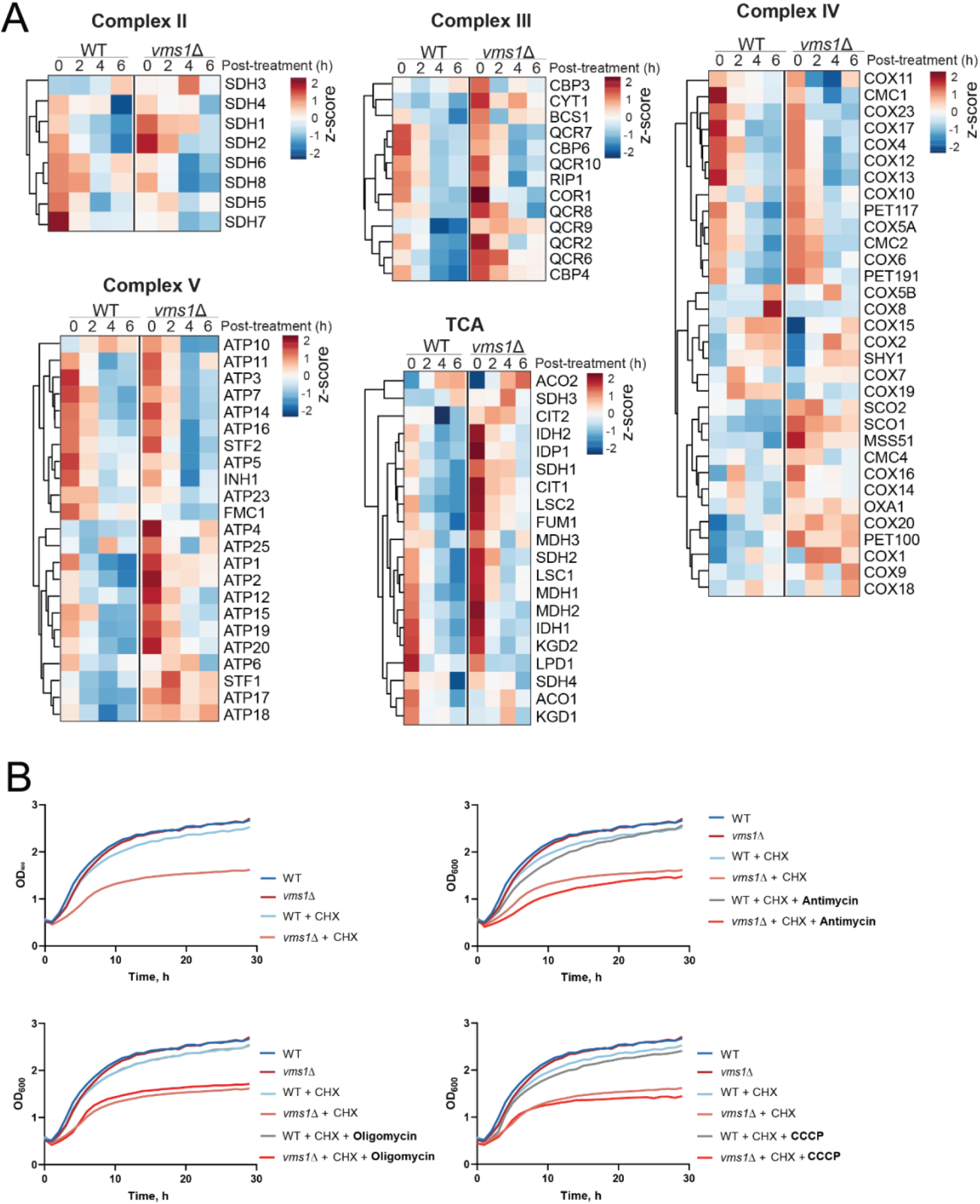
Loss of Vms1 alters the expression of mitochondrial respiratory-chain components but does not affect growth under mitochondrial stress. **(A)** Heatmaps showing the relative protein abundance of genes encoding components of respiratory-chain complexes II–V and enzymes of the tricarboxylic acid (TCA) cycle in WT and *vms1Δ* cells at 0, 2, 4, and 6 h after treatment. Values are row-normalized and displayed as *z*-scores. Proteins were hierarchically clustered according to their temporal expression profiles. **(B)** Growth of WT and *vms1Δ* cells in the absence or presence of 100 ng/ml CHX and the indicated mitochondrial inhibitors: 0.1 µg/ml antimycin, 0.1 µg/ml oligomycin, or 5 µM CCCP. Cells were diluted to an OD_600_ of 0.2, and growth was monitored by measuring OD_600_ over time.

**Figure S5.**
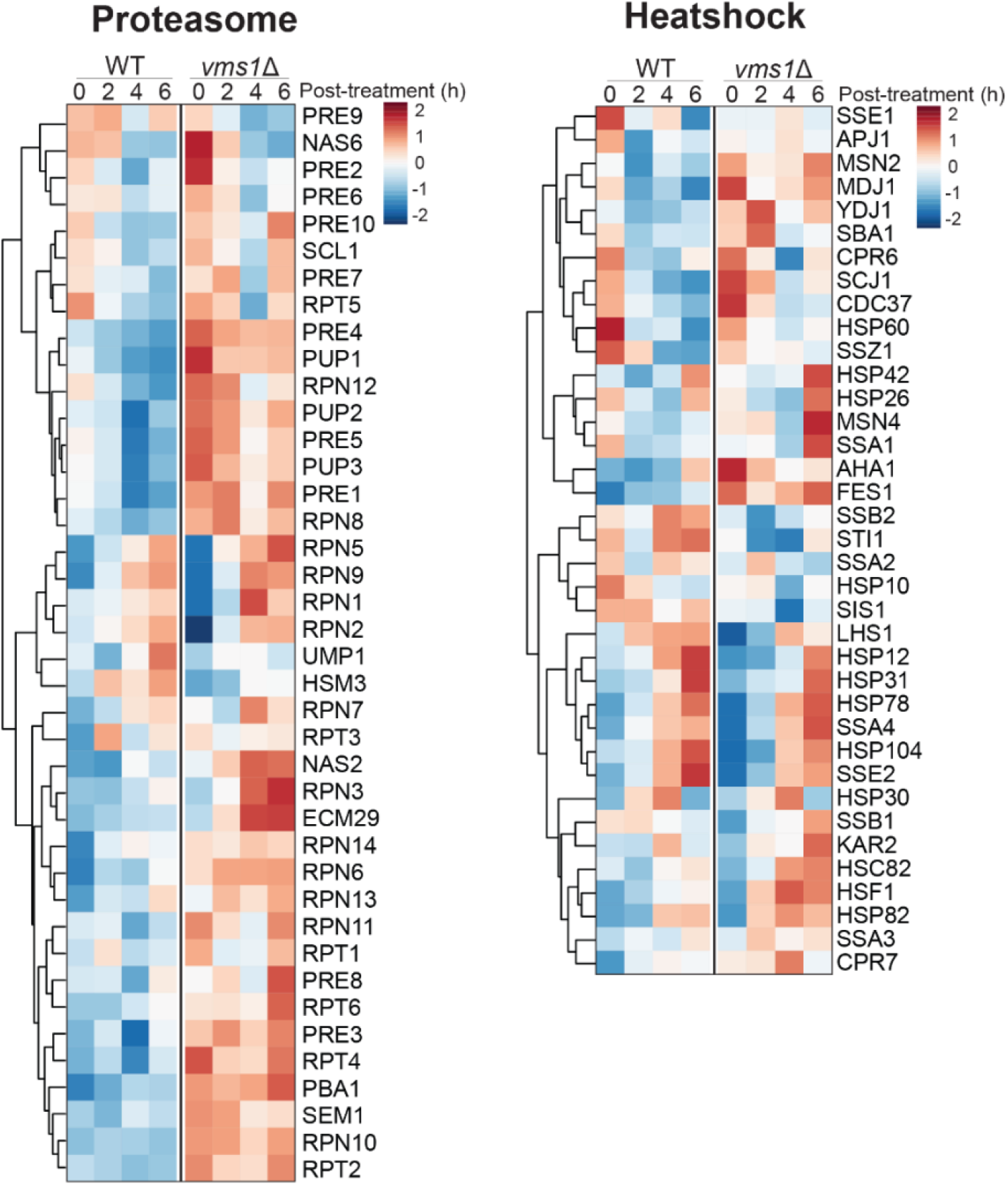
Loss of Vms1 induces heat-shock and proteasome-related responses. Heatmaps of heat-shock response- and proteasome-related proteins in WT and *vms1Δ* cells after 0-6 h of CHX treatment.

**Figure S6.**
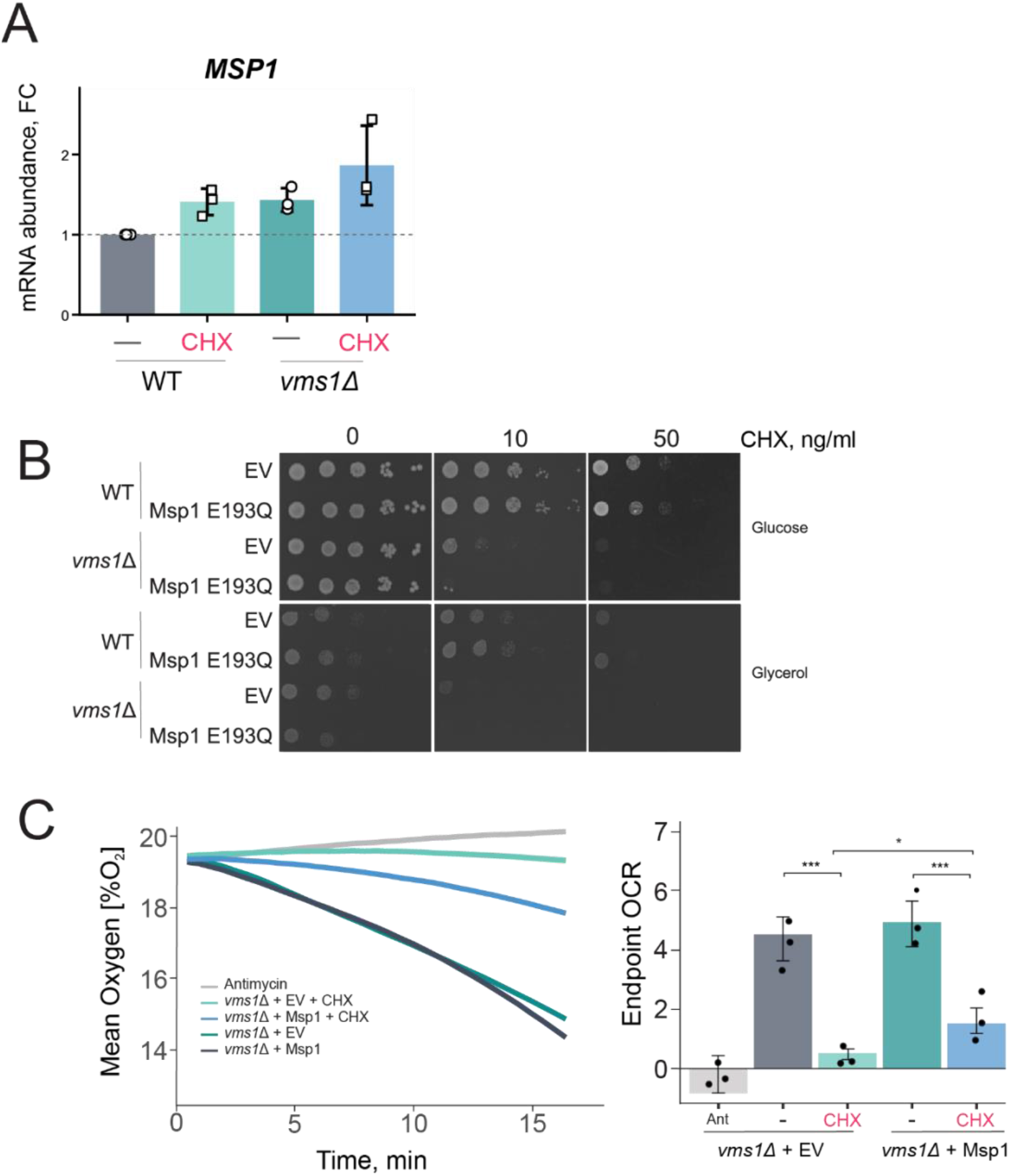
Msp1 expression partially restores respiration. **(A)** Relative MSP1 mRNA levels in WT and *vms1Δ* cells treated with or without 100 ng/ml CHX. Values are shown relative to untreated WT cells, and points represent biological replicates. Statistical significance was determined by two-way ANOVA with Tukey’s multiple-comparisons test. **(B)** The growth of WT and vms1Δ cells transformed with empty vector (EV) or a vector expressing ATPase-defective Msp1 E193Q mutant was analyzed by drop dilution assay on YPD or YPGlycerol with different CHX concentrations. **(C)** Oxygen consumption kinetics (left) and endpoint OCR values (right) of *vms1Δ* cells expressing WT Msp1 or carrying an empty vector (EV), with or without 100 ng/ml CHX. Antimycin-treated cells served as respiration-deficient control. Oxygen consumption was measured using FirePlate-O_2_ sensors and is expressed as percentage air saturation. Data represent three biological replicates. Statistical significance was determined by two-way ANOVA with Tukey’s multiple-comparisons test. *P < 0.05, **P < 0.01, ***P < 0.001, ****P < 0.0001.

**Figure S7.**
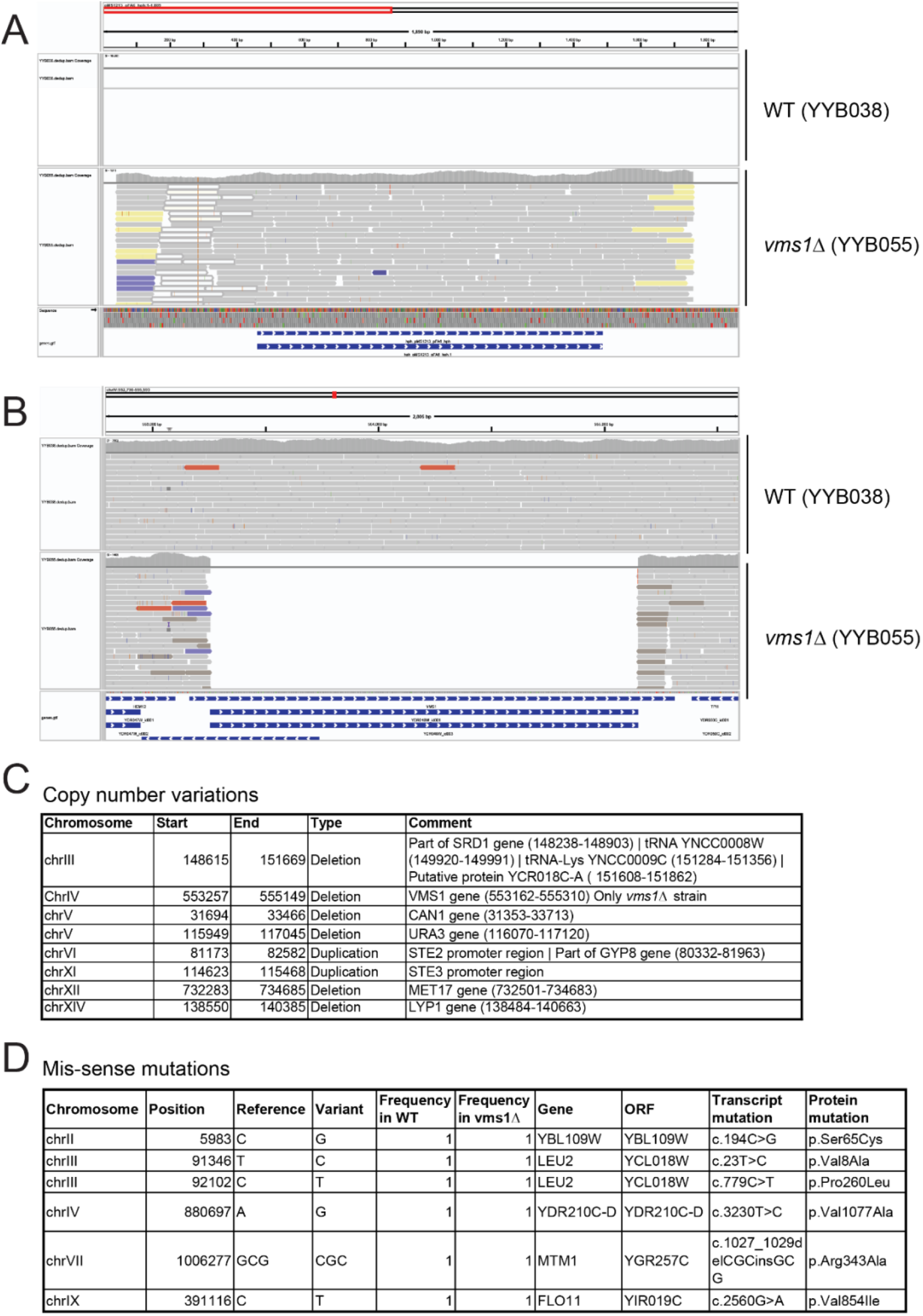
Verification of WT and *vms1Δ* strains by whole-genome sequencing. **(A)** Read alignments to the hygromycin resistance cassette sequence showing no reads in the WT strain (YYB038) and cassette integration in the *vms1Δ* strain. **(B)** Read alignments to the *VMS1* locus showing the presence of the gene and no copy number increase in the WT strain and complete loss of the gene in *vms1Δ* strain. **(C)** The summary of the copy number variations detected in WT and *vms1Δ* strains compared to the S288C reference genome with the only difference between WT and *vms1Δ* detected in VMS1 gene. **(D)** The summary of the mis-sense mutations in WT and *vms1Δ* strains compared to the S288C reference genome showing that there are no new mis-sense mutations in *vms1Δ*. See Methods for details.

## References

Banerjee, R., Trauschke, V., Bertram, N., Aretz, I., Osman, C., Lamb, D. C. and Mokranjac, D. (2026). mtHsp70 chaperone converts mitochondrial proteostasis stress into impaired protein import. Proceedings of the National Academy of Sciences 123, e2526136123.

Bengtson, M. H. and Joazeiro, C. A. P. (2010). Role of a ribosome-associated E3 ubiquitin ligase in protein quality control. Nature 467, 470–473.

Bertram, N., Izawa, T., Thoma, F., Schwenkert, S., Duvezin-Caubet, S., Park, S.-H., Wagener, N., Devin, A., Osman, C., Neupert, W., et al. (2025). Delayed protein translocation protects mitochondria against toxic CAT-tailed proteins. Molecular Cell 85, 4082–4092.e7.

Boos, F., Krämer, L., Groh, C., Jung, F., Haberkant, P., Stein, F., Wollweber, F., Gackstatter, A., Zöller, E., Laan, M. van der, et al. (2019). Mitochondrial protein-induced stress triggers a global adaptive transcriptional programme. Nat Cell Biol 21, 442–451.

Brandman, O., Stewart-Ornstein, J., Wong, D., Larson, A., Williams, C. C., Li, G.-W., Zhou, S., King, D., Shen, P. S., Weibezahn, J., et al. (2012). A Ribosome-Bound Quality Control Complex Triggers Degradation of Nascent Peptides and Signals Translation Stress. Cell 151, 1042–1054.

Breslow, D. K., Cameron, D. M., Collins, S. R., Schuldiner, M., Stewart-Ornstein, J., Newman, H. W., Braun, S., Madhani, H. D., Krogan, N. J. and Weissman, J. S. (2008). A comprehensive strategy enabling high-resolution functional analysis of the yeast genome. Nat Methods 5, 711–718.

Buechel, E. R. and Pinkett, H. W. (2020). Transcription factors and ABC transporters: from pleiotropic drug resistance to cellular signaling in yeast. FEBS Letters 594, 3943–3964.

Chen, S. (2025). fastp 1.0: An ultra-fast all-round tool for FASTQ data quality control and preprocessing. iMeta 4, e70078.

Chen, Y., Umanah, G. K. E., Dephoure, N., Andrabi, S. A., Gygi, S. P., Dawson, T. M., Dawson, V. L. and Rutter, J. (2014). Msp1/ATAD1 maintains mitochondrial function by facilitating the degradation of mislocalized tail-anchored proteins. EMBO J 33, 1548–1564.

Chiabudini, M., Tais, A., Zhang, Y., Hayashi, S., Wölfle, T., Fitzke, E. and Rospert, S. (2014). Release Factor eRF3 Mediates Premature Translation Termination on Polylysine-Stalled Ribosomes in Saccharomyces cerevisiae. Molecular and Cellular Biology 34, 4062–4076.

Choe, Y.-J., Park, S.-H., Hassemer, T., Körner, R., Vincenz-Donnelly, L., Hayer-Hartl, M. and Hartl, F. U. (2016). Failure of RQC machinery causes protein aggregation and proteotoxic stress. Nature 531, 191–195.

Chu, J., Hong, N. A., Masuda, C. A., Jenkins, B. V., Nelms, K. A., Goodnow, C. C., Glynne, R. J., Wu, H., Masliah, E., Joazeiro, C. A. P., et al. (2009). A mouse forward genetics screen identifies LISTERIN as an E3 ubiquitin ligase involved in neurodegeneration. Proceedings of the National Academy of Sciences 106, 2097–2103.

Couvillion, M. T., Soto, I. C., Shipkovenska, G. and Churchman, L. S. (2016). Synchronized mitochondrial and cytosolic translation programs. Nature 533, 499–503.

Danecek, P., Bonfield, J. K., Liddle, J., Marshall, J., Ohan, V., Pollard, M. O., Whitwham, A., Keane, T., McCarthy, S. A., Davies, R. M., et al. (2021). Twelve years of SAMtools and BCFtools. Gigascience 10, giab008.

Defenouillère, Q., Yao, Y., Mouaikel, J., Namane, A., Galopier, A., Decourty, L., Doyen, A., Malabat, C., Saveanu, C., Jacquier, A., et al. (2013). Cdc48-associated complex bound to 60S particles is required for the clearance of aberrant translation products. Proceedings of the National Academy of Sciences 110, 5046–5051.

Di Fraia, D., Marino, A., Lee, J. H., Kelmer Sacramento, E., Baumgart, M., Bagnoli, S., Balla, T., Schalk, F., Kamrad, S., Guan, R., et al. (2025). Altered translation elongation contributes to key hallmarks of aging in the killifish brain. Science 389, eadk3079.

D’Orazio, K. N., Wu, C. C.-C., Sinha, N., Loll-Krippleber, R., Brown, G. W. and Green, R. (2019). The endonuclease Cue2 cleaves mRNAs at stalled ribosomes during No Go Decay. eLife 8, e49117.

Engel, S. R., Aleksander, S., Nash, R. S., Wong, E. D., Weng, S., Miyasato, S. R., Sherlock, G. and Cherry, J. M. (2025). *Saccharomyces* Genome Database: advances in genome annotation, expanded biochemical pathways, and other key enhancements. GENETICS 229, iyae185.

Ennis, A., Wang, L., Xu, Y., Saidi, L., Wang, X., Yu, C., Yun, S., Huang, L. and Ye, Y. (2025). NEMF-mediated CAT tailing facilitates translocation-associated quality control. Journal of Cell Biology 224, e202408199.

Fer, E., Yao, T., McGrath, K. M., Goldman, A. D. and Kaçar, B. (2025). The origins and evolution of translation factors. Trends in Genetics 41, 590–600.

Fessler, E., Eckl, E.-M., Schmitt, S., Mancilla, I. A., Meyer-Bender, M. F., Hanf, M., Philippou-Massier, J., Krebs, S., Zischka, H. and Jae, L. T. (2020). A pathway coordinated by DELE1 relays mitochondrial stress to the cytosol. Nature 579, 433–437.

Filbeck, S., Cerullo, F., Pfeffer, S. and Joazeiro, C. A. P. (2022). Ribosome-associated quality-control mechanisms from bacteria to humans. Molecular Cell 82, 1451–1466.

Gale, A. N., Pavesic, M. W., Nickels, T. J., Xu, Z., Cormack, B. P. and Cunningham, K. W. (2023). Redefining pleiotropic drug resistance in a pathogenic yeast: Pdr1 functions as a sensor of cellular stresses in Candida glabrata. mSphere 8, e00254-23.

Garreau de Loubresse, N., Prokhorova, I., Holtkamp, W., Rodnina, M. V., Yusupova, G. and Yusupov, M. (2014). Structural basis for the inhibition of the eukaryotic ribosome. Nature 513, 517–522.

Garrison, E. and Marth, G. (2012). Haplotype-based variant detection from short-read sequencing.

Ge, S. X., Jung, D. and Yao, R. (2020). ShinyGO: a graphical gene-set enrichment tool for animals and plants. Bioinformatics 36, 2628–2629.

Giaever, G., Chu, A. M., Ni, L., Connelly, C., Riles, L., Véronneau, S., Dow, S., Lucau-Danila, A., Anderson, K., André, B., et al. (2002). Functional profiling of the *Saccharomyces cerevisiae* genome. Nature 418, 387–391.

Gietz, R. D. and Woods, R. A. (2006). Yeast Transformation by the LiAc/SS Carrier DNA/PEG Method. In Yeast Protocol (ed. Xiao, W.), pp. 107–120. Totowa, NJ: Humana Press.

Guo, X., Aviles, G., Liu, Y., Tian, R., Unger, B. A., Lin, Y.-H. T., Wiita, A. P., Xu, K., Correia, M. A. and Kampmann, M. (2020). Mitochondrial stress is relayed to the cytosol by an OMA1–DELE1–HRI pathway. Nature 579, 427–432.

Guydosh, N. R. and Green, R. (2014). Dom34 Rescues Ribosomes in 3′ Untranslated Regions. Cell 156, 950–962.

Hanscho, M., Ruckerbauer, D. E., Chauhan, N., Hofbauer, H. F., Krahulec, S., Nidetzky, B., Kohlwein, S. D., Zanghellini, J. and Natter, K. (2012). Nutritional requirements of the BY series of Saccharomyces cerevisiae strains for optimum growth. FEMS Yeast Research 12, 796–808.

Hansen, K. G., Aviram, N., Laborenz, J., Bibi, C., Meyer, M., Spang, A., Schuldiner, M. and Herrmann, J. M. (2018). An ER surface retrieval pathway safeguards the import of mitochondrial membrane proteins in yeast. Science 361, 1118–1122.

Hoseini, H., Pandey, S., Jores, T., Schmitt, A., Franz-Wachtel, M., Macek, B., Buchner, J., Dimmer, K. S. and Rapaport, D. (2016). The cytosolic cochaperone Sti1 is relevant for mitochondrial biogenesis and morphology. The FEBS Journal 283, 3338–3352.

Ikeuchi, K., Tesina, P., Matsuo, Y., Sugiyama, T., Cheng, J., Saeki, Y., Tanaka, K., Becker, T., Beckmann, R. and Inada, T. (2019). Collided ribosomes form a unique structural interface to induce Hel2-driven quality control pathways. The EMBO Journal 38, e100276.

Inada, T. (2026). Ribosome-associated quality control and related mechanisms. Nat Struct Mol Biol 33, 394–407.

Ishimura, R., Nagy, G., Dotu, I., Zhou, H., Yang, X.-L., Schimmel, P., Senju, S., Nishimura, Y., Chuang, J. H. and Ackerman, S. L. (2014). Ribosome stalling induced by mutation of a CNS-specific tRNA causes neurodegeneration. Science 345, 455–459.

Iyer, K. V., Walter, C. A., Kraft, A.-A., Müller, M., Tittel, L. S. and Winz, M.-L. (2025). Jlp2 is an RQC complex-independent release factor acting on aberrant peptidyl-tRNA, protecting cells against translation elongation stress. 2025.09.04.673968.

Izawa, T., Tsuboi, T., Kuroha, K., Inada, T., Nishikawa, S. and Endo, T. (2012). Roles of Dom34:Hbs1 in Nonstop Protein Clearance from Translocators for Normal Organelle Protein Influx. Cell Reports 2, 447–453.

Izawa, T., Park, S.-H., Zhao, L., Hartl, F. U. and Neupert, W. (2017). Cytosolic Protein Vms1 Links Ribosome Quality Control to Mitochondrial and Cellular Homeostasis. Cell 171, 890–903.e18.

Janke, C., Magiera, M. M., Rathfelder, N., Taxis, C., Reber, S., Maekawa, H., Moreno-Borchart, A., Doenges, G., Schwob, E., Schiebel, E., et al. (2004). A versatile toolbox for PCR-based tagging of yeast genes: new fluorescent proteins, more markers and promoter substitution cassettes. Yeast 21, 947–962.

Juszkiewicz, S., Chandrasekaran, V., Lin, Z., Kraatz, S., Ramakrishnan, V. and Hegde, R. S. (2018). ZNF598 Is a Quality Control Sensor of Collided Ribosomes. Molecular Cell 72, 469–481.e7.

Juszkiewicz, S., Peak-Chew, S.-Y. and Hegde, R. S. (2025). Mechanism of chaperone recruitment and retention on mitochondrial precursors. MBoC 36, ar39.

Kim, J., Goldstein, M., Zecchel, L., Ghorayeb, R., Maxwell, C. A. and Weidberg, H. (2024). ATAD1 prevents clogging of TOM and damage caused by un-imported mitochondrial proteins. Cell Reports 43, 114473.

Kostova, K. K., Hickey, K. L., Osuna, B. A., Hussmann, J. A., Frost, A., Weinberg, D. E. and Weissman, J. S. (2017). CAT-tailing as a fail-safe mechanism for efficient degradation of stalled nascent polypeptides. Science 357, 414–417.

Kuroha, K., Zinoviev, A., Hellen, C. U. T. and Pestova, T. V. (2018). Release of Ubiquitinated and Non-ubiquitinated Nascent Chains from Stalled Mammalian Ribosomal Complexes by ANKZF1 and Ptrh1. Molecular Cell 72, 286–302.e8.

Li, H. (2013). Aligning sequence reads, clone sequences and assembly contigs with BWA-MEM.

Li, W., Scheel, T. and Shen, P. S. (2025a). Mechanism of nascent chain removal by the ribosome-associated quality control complex. Nat Commun 16, 5792.

Li, Y., Liu, Y., Jiang, Y., Yang, Y., Ni, W., Zhang, W. and Tan, L. (2025b). New antifungal strategies and drug development against WHO critical priority fungal pathogens. Front Cell Infect Microbiol 15, 1662442.

Livak, K. J. and Schmittgen, T. D. (2001). Analysis of relative gene expression data using real-time quantitative PCR and the 2(-Delta Delta C(T)) Method. Methods 25, 402–408.

Longtine, M. S., Iii, A. M., Demarini, D. J., Shah, N. G., Wach, A., Brachat, A., Philippsen, P. and Pringle, J. R. (1998). Additional modules for versatile and economical PCR-based gene deletion and modification in Saccharomyces cerevisiae. Yeast 14, 953–961.

Lyumkis, D., Oliveira dos Passos, D., Tahara, E. B., Webb, K., Bennett, E. J., Vinterbo, S., Potter, C. S., Carragher, B. and Joazeiro, C. A. P. (2014). Structural basis for translational surveillance by the large ribosomal subunit-associated protein quality control complex. Proceedings of the National Academy of Sciences 111, 15981–15986.

Mårtensson, C. U., Priesnitz, C., Song, J., Ellenrieder, L., Doan, K. N., Boos, F., Floerchinger, A., Zufall, N., Oeljeklaus, S., Warscheid, B., et al. (2019). Mitochondrial protein translocation-associated degradation. Nature 569, 679–683.

Martin, P. B., Kigoshi-Tansho, Y., Sher, R. B., Ravenscroft, G., Stauffer, J. E., Kumar, R., Yonashiro, R., Müller, T., Griffith, C., Allen, W., et al. (2020). NEMF mutations that impair ribosome-associated quality control are associated with neuromuscular disease. Nat Commun 11, 4625.

Matsuo, Y., Ikeuchi, K., Saeki, Y., Iwasaki, S., Schmidt, C., Udagawa, T., Sato, F., Tsuchiya, H., Becker, T., Tanaka, K., et al. (2017). Ubiquitination of stalled ribosome triggers ribosome-associated quality control. Nat Commun 8, 159.

Metzl-Raz, E., Kafri, M., Yaakov, G., Soifer, I., Gurvich, Y. and Barkai, N. (2017). Principles of cellular resource allocation revealed by condition-dependent proteome profiling. eLife 6, e28034.

Nanjaraj Urs, A. N., Lasehinde, V., Kim, L., McDonald, E., Yan, L. L. and Zaher, H. S. (2024). Inability to rescue stalled ribosomes results in overactivation of the integrated stress response. Journal of Biological Chemistry 300, 107290.

Neupert, W. and Herrmann, J. M. (2007). Translocation of Proteins into Mitochondria. Annual Review of Biochemistry 76, 723–749.

Opaliński, Ł., Song, J., Priesnitz, C., Wenz, L.-S., Oeljeklaus, S., Warscheid, B., Pfanner, N. and Becker, T. (2018). Recruitment of Cytosolic J-Proteins by TOM Receptors Promotes Mitochondrial Protein Biogenesis. Cell Reports 25, 2036–2043.e5.

Pochopien, A. A., Beckert, B., Kasvandik, S., Berninghausen, O., Beckmann, R., Tenson, T. and Wilson, D. N. (2021). Structure of Gcn1 bound to stalled and colliding 80S ribosomes. Proceedings of the National Academy of Sciences 118, e2022756118.

R Core Team (2020). R: A language and environment for statistical computing. R Foundation for Statistical Computing, Vienna, Austria.

Rausch, T., Zichner, T., Schlattl, A., Stütz, A. M., Benes, V. and Korbel, J. O. (2012). DELLY: structural variant discovery by integrated paired-end and split-read analysis. Bioinformatics 28, i333–i339.

Robinson, J. T., Thorvaldsdóttir, H., Winckler, W., Guttman, M., Lander, E. S., Getz, G. and Mesirov, J. P. (2011). Integrative genomics viewer. Nat Biotechnol 29, 24–26.

Schuller, A. P. and Green, R. (2018). Roadblocks and resolutions in eukaryotic translation. Nat Rev Mol Cell Biol 19, 526–541.

Schulte, U., den Brave, F., Haupt, A., Gupta, A., Song, J., Müller, C. S., Engelke, J., Mishra, S., Mårtensson, C., Ellenrieder, L., et al. (2023). Mitochondrial complexome reveals quality-control pathways of protein import. Nature 614, 153–159.

Shao, S., Brown, A., Santhanam, B. and Hegde, R. S. (2015). Structure and Assembly Pathway of the Ribosome Quality Control Complex. Molecular Cell 57, 433–444.

Shcherbik, N., Chernova, T. A., Chernoff, Y. O. and Pestov, D. G. (2016). Distinct types of translation termination generate substrates for ribosome-associated quality control. Nucleic Acids Res 44, 6840–6852.

Shen, P. S., Park, J., Qin, Y., Li, X., Parsawar, K., Larson, M. H., Cox, J., Cheng, Y., Lambowitz, A. M., Weissman, J. S., et al. (2015). Rqc2p and 60S ribosomal subunits mediate mRNA-independent elongation of nascent chains. Science 347, 75–78.

Simms, C. L., Yan, L. L. and Zaher, H. S. (2017). Ribosome Collision Is Critical for Quality Control during No-Go Decay. Molecular Cell 68, 361–373.e5.

Sinha, N. K., McKenney, C., Yeow, Z. Y., Li, J. J., Nam, K. H., Yaron-Barir, T. M., Johnson, J. L., Huntsman, E. M., Cantley, L. C., Ordureau, A., et al. (2024). The ribotoxic stress response drives UV-mediated cell death. Cell 187, 3652–3670.e40.

Sitron, C. S. and Brandman, O. (2019). CAT tails drive degradation of stalled polypeptides on and off the ribosome. Nat Struct Mol Biol 26, 450–459.

Sitron, C. S., Park, J. H., Giafaglione, J. M. and Brandman, O. (2020). Aggregation of CAT tails blocks their degradation and causes proteotoxicity in S. cerevisiae. PLOS ONE 15, e0227841.

Su, T., Izawa, T., Thoms, M., Yamashita, Y., Cheng, J., Berninghausen, O., Hartl, F. U., Inada, T., Neupert, W. and Beckmann, R. (2019). Structure and function of Vms1 and Arb1 in RQC and mitochondrial proteome homeostasis. Nature 1.

Tanenbaum, M. E., Gilbert, L. A., Qi, L. S., Weissman, J. S. and Vale, R. D. (2014). A protein tagging system for signal amplification in gene expression and fluorescence imaging. Cell 159, 635–646.

Udagawa, T., Seki, M., Okuyama, T., Adachi, S., Natsume, T., Noguchi, T., Matsuzawa, A. and Inada, T. (2021). Failure to Degrade CAT-Tailed Proteins Disrupts Neuronal Morphogenesis and Cell Survival. Cell Reports 34, 108599.

Vallières, C., Raulo, R., Dickinson, M. and Avery, S. V. (2018). Novel Combinations of Agents Targeting Translation That Synergistically Inhibit Fungal Pathogens. Front. Microbiol. 9,.

van den Elzen, A. M. G., Schuller, A., Green, R. and Séraphin, B. (2014). Dom34-Hbs1 mediated dissociation of inactive 80S ribosomes promotes restart of translation after stress. The EMBO Journal 33, 265–276.

van Haaften-Visser, D. Y., Harakalova, M., Mocholi, E., van Montfrans, J. M., Elkadri, A., Rieter, E., Fiedler, K., van Hasselt, P. M., Triffaux, E. M. M., van Haelst, M. M., et al. (2017). Ankyrin repeat and zinc-finger domain-containing 1 mutations are associated with infantile-onset inflammatory bowel disease. Journal of Biological Chemistry 292, 7904–7920.

Verma, R., Reichermeier, K. M., Burroughs, A. M., Oania, R. S., Reitsma, J. M., Aravind, L. and Deshaies, R. J. (2018). Vms1 and ANKZF1 peptidyl-tRNA hydrolases release nascent chains from stalled ribosomes. Nature 557, 446–451.

Wagih, O., Usaj, M., Baryshnikova, A., VanderSluis, B., Kuzmin, E., Costanzo, M., Myers, C. L., Andrews, B. J., Boone, C. M. and Parts, L. (2013). SGAtools: one-stop analysis and visualization of array-based genetic interaction screens. Nucleic Acids Res 41, W591–W596.

Wang, X. and Chen, X. J. (2015). A cytosolic network suppressing mitochondria-mediated proteostatic stress and cell death. Nature 524, 481–484.

Weidberg, H. and Amon, A. (2018). MitoCPR—A surveillance pathway that protects mitochondria in response to protein import stress. Science 360, eaan4146.

Wohlever, M. L., Mateja, A., McGilvray, P. T., Day, K. J. and Keenan, R. J. (2017). Msp1 Is a Membrane Protein Dislocase for Tail-Anchored Proteins. Molecular Cell 67, 194–202.e6.

Wrobel, L., Topf, U., Bragoszewski, P., Wiese, S., Sztolsztener, M. E., Oeljeklaus, S., Varabyova, A., Lirski, M., Chroscicki, P., Mroczek, S., et al. (2015). Mistargeted mitochondrial proteins activate a proteostatic response in the cytosol. Nature 524, 485–488.

Wu, C. C.-C., Peterson, A., Zinshteyn, B., Regot, S. and Green, R. (2020). Ribosome Collisions Trigger General Stress Responses to Regulate Cell Fate. Cell 182, 404–416.e14.

Yan, L. L. and Zaher, H. S. (2021). Ribosome quality control antagonizes the activation of the integrated stress response on colliding ribosomes. Molecular Cell 81, 614–628.e4.

Yan, L. L., Simms, C. L., McLoughlin, F., Vierstra, R. D. and Zaher, H. S. (2019). Oxidation and alkylation stresses activate ribosome-quality control. Nat Commun 10, 5611.

Yofe, I. and Schuldiner, M. (2014). Primers-4-Yeast: a comprehensive web tool for planning primers for Saccharomyces cerevisiae. Yeast 31, 77–80.

Yonashiro, R., Tahara, E. B., Bengtson, M. H., Khokhrina, M., Lorenz, H., Chen, K.-C., Kigoshi-Tansho, Y., Savas, J. N., Yates, J. R.III, Kay, S. A., et al. (2016). The Rqc2/Tae2 subunit of the ribosome-associated quality control (RQC) complex marks ribosome-stalled nascent polypeptide chains for aggregation. eLife 5, e11794.

Young, J. C., Hoogenraad, N. J. and Hartl, F. U. (2003). Molecular Chaperones Hsp90 and Hsp70 Deliver Preproteins to the Mitochondrial Import Receptor Tom70. Cell 112, 41–50.

Yuan, Z., Balzarini, M., Volpe, M., Goldstein, M., Peng, T. S., Hui, E., Fang, N. N., Albihlal, W. S., Hajimohammadi, M., Wei, K., et al. (2026). A direct role for a mitochondrial targeting sequence in signalling stress. Nature 649, 1302–1311.

Zara, V., Ferramosca, A., Robitaille-Foucher, P., Palmieri, F. and Young, J. C. (2009). Mitochondrial carrier protein biogenesis: role of the chaperones Hsc70 and Hsp90. Biochemical Journal 419, 369–375.

Zhou, C., Zhang, M., Murray, J., Paulo, J., Gygi, S., Shao, S., Whitman, M. and Keller, T. (2025). GCN1 couples GCN2 to ribosomal state to initiate amino acid response pathway signaling. Science 390, eads8728.

Zhu, L., Wu, K., You, J., Mi, W., Xu, J., Li, L., Yang, F., Xia, X., Yan, H., Li, F., et al. (2026). Mitochondrial glutamine import sustains electron transport chain integrity independently of glutaminolysis in cancer. Molecular Cell 86, 150–165.e9.

Zurita Rendón, O., Fredrickson, E. K., Howard, C. J., Van Vranken, J., Fogarty, S., Tolley, N. D., Kalia, R., Osuna, B. A., Shen, P. S., Hill, C. P., et al. (2018). Vms1p is a release factor for the ribosome-associated quality control complex. Nat Commun 9, 2197.

